# *Drosophila* HS dendrites are resilient to adult-onset deficits in mitochondrial dynamics

**DOI:** 10.1101/2025.11.07.687201

**Authors:** Hailey Q. Wang, Demetria G. Fortson, Eavan D. Burton, Hunter S. Whitbeck, Briana M. Robles, Letitia Bortey, Avi J. Adler, Jordan I. Kalai, Giovanna Napoleone, Erin L. Barnhart

## Abstract

Mitochondrial transport, fusion, and fission are necessary for neuronal development, but the role of mitochondrial dynamics in neuronal maintenance remains unclear. In this work, we employed functional *in vivo* imaging of neurons in the *Drosophila* visual system, HS (“horizontal system”) cells, to determine how adult-onset deficits in mitochondrial dynamics affect mitochondrial localization, local regulation of ATP, and dendrite maintenance. In mature HS neurons, inhibition of mitochondrial transport or fusion depleted mitochondria from the dendrite over time but, surprisingly, had no effect on dendrite morphology. Moreover, adult-restricted mitochondrial mis- localization affected neither visual stimulus-driven dendritic calcium responses nor local, dynamic regulation of ATP levels. In contrast, when induced during development, the same perturbations caused mitochondrial mis-localization, loss of dendrite complexity, abrogation of stimulus-locked calcium responses and ATP fluctuations, and age-dependent dendrite degeneration. Thus, although mitochondrial dynamics are necessary during neuronal development, mature dendrites are capable of maintaining form and function *in vivo* in the absence of properly-positioned mitochondria.

## INTRODUCTION

Power-hungry neurons are widely believed to rely on proper localization of healthy mitochondria to coordinate ATP production with fluctuating energetic demands (Devine and Kittler, 2018; Misgeld and Schwarz, 2017; Sheng, 2017). Maintenance of properly-localized, healthy mitochondria, or mitostasis, is particularly challenging in neurons (Misgeld and Schwarz, 2017). Mitochondria and their constituent elements (e.g. mitochondrial proteins) have substantially shorter lifespans than neurons (Vincow et al., 2013), and the vast majority of mitochondrial proteins are encoded by the nuclear genome. Therefore, neurons are thought to rely on mitochondrial motility to distribute mitochondria supplied with newly-synthesized proteins to axonal and dendritic arbors far-removed from the cell soma (Misgeld and Schwarz, 2017). Mitochondrial fusion and fission, in turn, are thought to contribute to mitostasis by facilitating the equitable distribution of functional mitochondrial components throughout the mitochondrial network (via fusion-dependent exchange of mitochondrial contents) as well as selective degradation of damaged mitochondria (via fission-dependent segregation) (Chan, 2020). According to this model, adult-onset deficits in mitochondrial dynamics ought to disrupt mitostasis and thus local production of ATP in neurites, ultimately resulting in neurodegeneration. However, although there is abundant evidence that mitochondrial motility, fusion, and fission are necessary for neuronal development (Courchet et al., 2013; Dickey and Strack, 2011; Dubal et al., 2022; Fukumitsu et al., 2015; Ishihara et al., 2009; Iwata et al., 2020; Kochan et al., 2024; Li et al., 2004; Lopez-Domenech et al., 2016; Morris and Hollenbeck, 1993; Troger et al., 2025), the extent to which mitochondrial dynamics contribute to maintenance of fully mature neurons *in vivo* remains unknown.

The role of mitochondrial dynamics in neuronal maintenance is poorly defined for several reasons. First, it is unclear how much mitochondrial transport, fusion, and fission occurs in fully mature neurons *in vivo*. Mitochondrial transport has been measured in neurons *in vivo* (Donovan et al., 2024; Mandal et al., 2021; Plucinska et al., 2012; Silva et al., 2021; Vagnoni and Bullock, 2018), but in some cell types transport declines dramatically as neurons mature (Faits et al., 2016; Lewis et al., 2016; Smit-Rigter et al., 2016), suggesting that mitochondrial motility may be more important for neuronal development than for neuronal maintenance. Moreover, although mitochondrial fusion and fission have been observed in neurons *ex vivo* (Fenton et al., 2024; Kochan et al., 2024; Lewis et al., 2018), fusion and fission have yet to be measured in neurons *in vivo*. Second, deficits in mitochondrial transport, fusion, and fission reduce neuronal morphological complexity (Berthet et al., 2014; Courchet et al., 2013; Dickey and Strack, 2011; Fukumitsu et al., 2015; Ishihara et al., 2009; Lopez-Domenech et al., 2016; Richhariya et al., 2025) and disrupt synaptic plasticity (Divakaruni et al., 2018; Li et al., 2004) and synaptic transmission (Lewis et al., 2018; Verstreken et al., 2005; Vevea and Chapman, 2023). However, with limited exceptions (Berthet et al., 2014; Lopez-Domenech et al., 2016), these mitochondrial deficits have been examined either in cell culture or in the context of genetic manipulations that affect neuronal development and/or large populations of neurons *in vivo*, making it difficult to determine whether neuronal dysfunction is due to cell-intrinsic defects in mitostasis that disrupt neuronal maintenance versus developmental effects that disrupt neuronal growth. Indeed, when induced in mid-life, pan-neuronal inhibition of mitochondrial fusion or fission, rather than causing neuronal degeneration, results in lifespan extension in *Drosophila* (Rana et al., 2017). Lastly, experiments in cultured neurons have shown that mitochondrial oxidative phosphorylation (OXPHOS) is necessary for activity-dependent upregulation of ATP production (Rangaraju et al., 2014), but there is no direct evidence linking cell type-specific deficits in mitochondrial transport, fusion, or fission to disrupted energy homeostasis in neurons *in vivo*. Altogether, it is necessary to define the cell-intrinsic effects of disrupted mitochondrial dynamics on neuronal maintenance.

In this work, we employed *Drosophila* genetic tools and functional *in vivo* imaging to measure and manipulate mitochondrial dynamics in a cell type-specific and temporally-restricted fashion. We focused in particular on neurons in the fly visual system called HS (“horizontal system”) cells. There are three HS neurons per optic lobe, each of which has an elaborately branched dendrite that innervates the third neuropil of the visual system, the lobula plate (Figure 1A-B). HS dendrites are densely packed with mitochondria (Figure 1B) (Donovan et al., 2024), and their morphologies and functional properties are well-defined (Barnhart et al., 2018; Busch et al., 2018; Cuntz et al., 2008; Erginkaya et al., 2025; Kim et al., 2017; Krapp et al., 1998; Schnell et al., 2010; Scott et al., 2002). In addition, we demonstrate here that visual input drives sustained increases in ATP levels in HS dendrites, making them ideal for interrogating the link between mitochondrial dynamics, local ATP production, and the maintenance of neuronal form and function. By measuring mitochondrial motility and content exchange in HS dendrites in the adult, we show that mitochondria in fully developed HS neurons are in fact dynamic. We further demonstrate that inhibiting mitochondrial transport or fusion in HS neurons during development causes mitochondrial mis-localization and/or fragmentation, reduced dendrite complexity, and age-dependent dendrite degeneration. In contrast, when restricted to the adult, the same perturbations cause depletion of mitochondria from the dendrite over time, but have no measurable effects on dendrite morphology or stimulus-driven ATP responses. Thus, HS dendrites are surprisingly resilient to adult-onset deficits in mitochondrial dynamics and localization.

**Figure 1:**
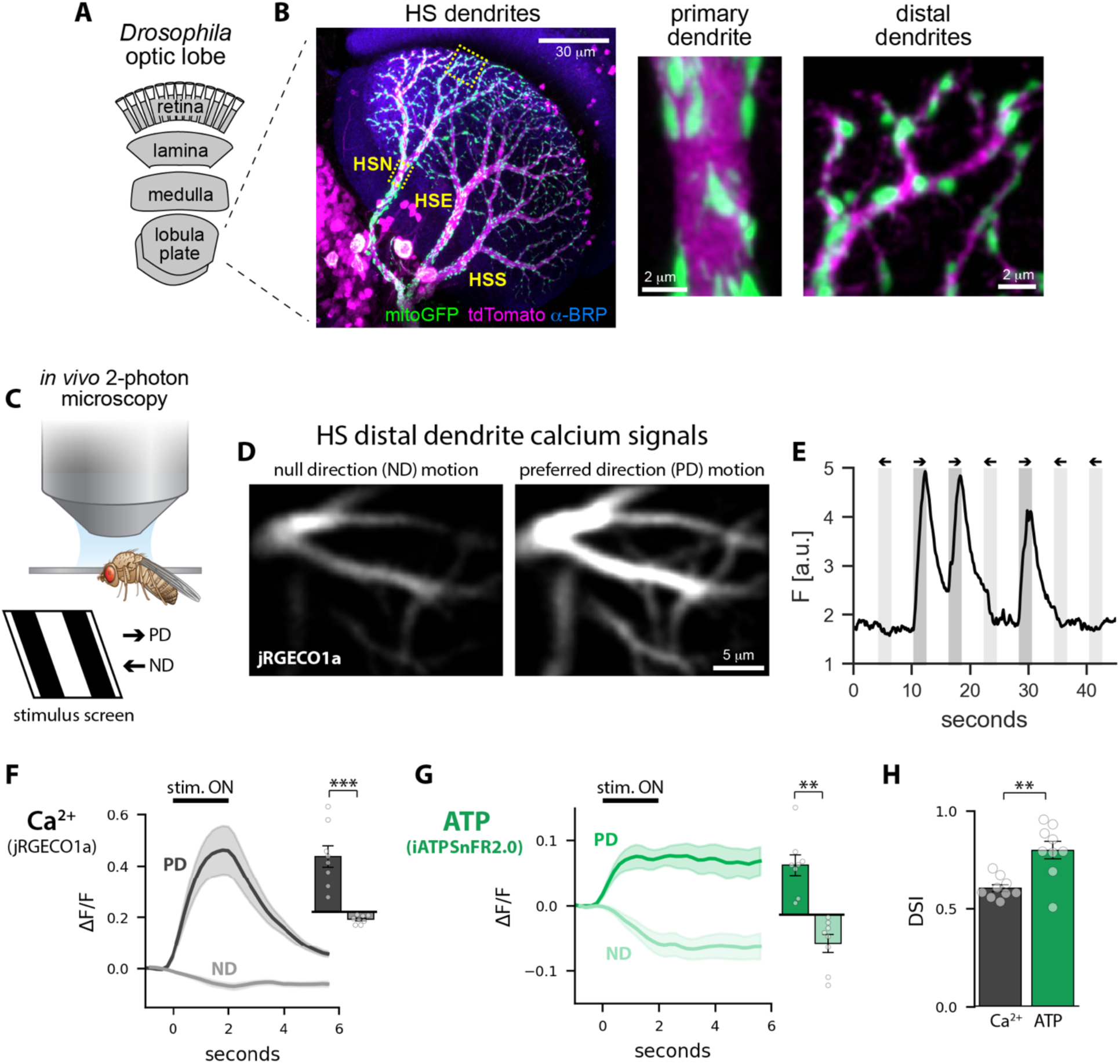
Visual input drives ATP responses in HS dendrites. A: Organization of the *Drosophila* visual system. B: Representative image of the dendrites of three HS neurons in the optic lobe (HSN, HSE, and HSS). The neurons were labeled with a cytosolic volume marker (tdTomato, magenta) and a mitochondria-targeted GFP (mitoGFP, green); the neuropil was labelled by immunostaining for the synaptic marker BRP (α-BRP, blue). Dashed yellow boxes indicate the primary and distal dendrites shown on the right. All images are maximum projections of z-stacks acquired using confocal Airyscan imaging. C: Experimental setup. *In vivo* two photon imaging of HS dendrites in a head-fixed fly with simultaneous visual stimulus presentation. The visual stimulus — full contrast square wave gratings that spanned the entire stimulus screen — moved in either the preferred direction (PD) for HS neurons (front-to-back, with respect to the fly’s eye) or in the opposite, null direction (ND). D: Two-photon images (average projections of a time series) of cytosolic calcium (jRGECO1a) in HS distal dendrites during presentation of gratings moving in the null direction (ND, left image) or preferred direction (PD, right image). E: Example jRGECO signals, plotted over time, in response to moving gratings. Shaded regions on the graph indicate when the grating was moving; light gray indicates ND motion and dark gray indicates PD motion. Two second bouts of motion were interspersed with four second presentations of static gratings. F-G: Average calcium (jRGECO1a, F) and ATP (iATPSnFR2.0, G) responses to ND and PD motion, plotted over time. Insets show average response amplitudes. H: Direction selectivity index (DSI = ½ (Rpref-Rnull)/Rpref) for calcium (gray) and ATP responses (green). Shading on line plots and error bars on bar plots indicate standard error of the mean; dots overlaid on the box plots indicate average response amplitudes for individual flies (N = 9 flies). Asterisks indicate significant differences (** p < 0.01, *** p < 0.001, paired T test).

## RESULTS

### Visual input drives ATP responses in HS dendrites

To investigate how deficits in mitostasis affect local, dynamic regulation of ATP levels, we first set out to measure visual stimulus-driven ATP fluctuations in HS dendrites. In cell culture, neurons upregulate ATP production in an activity-dependent fashion (Rangaraju et al., 2014). In *Drosophila*, stochastic fluctuations in neuronal calcium signals correlate with fluctuations in ATP across brain regions, suggesting that neuronal activity also upregulates ATP production *in vivo* (Mann et al., 2021). HS neurons are global motion detectors that selectively respond to specific patterns of global optic flow evoked by self-motion of the fly. Specifically, HS neurons depolarize in response to a preferred direction (PD) of visual global motion (front-to-back with respect to the fly’s eye) and hyperpolarize in response to global motion in the opposite, null direction (ND, back-to-front with respect to the fly’s eye) (Schnell et al., 2010). HS direction-selectivity is mediated by excitatory and inhibitory input onto HS dendrites: local motion detectors activated by front-to-back motion provide direct excitatory input while inhibitory interneurons relay input from neurons activated by local motion in the opposite direction (Maisak et al., 2013; Mauss et al., 2015; Nern et al., 2025). To determine how global motion affects ATP levels in HS dendrites, we used the GAL4/UAS system to express genetically encoded fluorescent reporters for ATP (iATPSnFR2.0 (Marvin et al., 2024)) and calcium (jRGECO1a (Dana et al., 2016)) in HS neurons. We used a GAL4 driver line that drives detectable expression in HS neurons in late pupal stages as well as throughout adulthood, with expression limited to HS neurons in the optic lobe (Figure S1). To measure stimulus-driven calcium and ATP responses, we used *in vivo* two photon microscopy to image HS dendrites in head-fixed flies while simultaneously projecting visual stimuli on a screen positioned in front of the fly (Figure 1C). We presented full contrast square wave gratings that spanned the entire stimulus screen (∼60x60degrees) and moved in either the preferred or null direction. Two second bouts of global motion were interspersed with four second periods without motion (i.e. presentation of a static grating), and bouts of PD versus ND motion were presented in a randomized order. As expected based on previously published work (Barnhart et al., 2018), we measured direction-selective calcium responses to global motion: PD motion drove large increases in jRGECO1a fluorescence whereas ND motion triggered a slight reduction in fluorescence (Figure 1D-F).

Strikingly, we also measured stimulus-locked fluctuations in ATP levels (Figure 1G). PD motion drove a sustained increase in iATPSnFR2.0 fluorescence, suggesting excitatory input onto HS dendrites triggers upregulation of ATP synthesis, with ATP production outpacing any stimulus-driven increase in ATP consumption. In contrast, ND motion caused a sustained reduction in fluorescence, suggesting inhibitory input increases ATP consumption without triggering a commensurate increase in ATP production. We quantified the direction selectivity of the calcium and ATP responses using a direction selectivity index (DSI) in which a value of 1 indicates perfect direction selectivity and a value of 0 indicates no direction selectivity (see Methods). ATP responses were significantly more direction selective than calcium responses due to the roughly equivalent absolute amplitudes of ATP increases versus decreases in response to PD versus ND motion (Figure 1H). Our ability to measure stimulus-locked fluctuations in ATP levels makes HS dendrites an ideal model for investigating how deficits in mitostasis affect dynamic regulation of ATP in a physiological context.

### Mitochondrial transport and content exchange in HS dendrites

Having established that visual input drives fluctuations in ATP in HS dendrites *in vivo*, we next wished to quantify mitochondrial dynamics in the same system. In previous work, we measured substantial mitochondrial transport in HS dendrites in young flies (Donovan et al., 2024). However, in some cell types, mitochondrial transport declines to nearly undetectable levels as neurons mature (Faits et al., 2016; Lewis et al., 2016; Smit-Rigter et al., 2016), casting some doubt on the extent to which mitochondrial transport contributes to mitostasis in adult neurons. To determine whether mitochondrial transport in HS dendrites continues throughout adulthood, we expressed a mitochondria-targeted GFP (mitoGFP) and a cytosolic volume marker (tdTomato) in HS neurons and we used *in vivo* confocal microscopy to image motile mitochondria in head-fixed flies ranging from two days to two months old (Figure 2A-H). To resolve individual motile mitochondria, we photobleached stationary mitochondria in the primary dendrite before collecting time lapse images (Figure 2B-D). Consistent with our previous work (Donovan et al., 2024), we observed mitochondria moving through the primary dendrite in both the anterograde direction (away from the cell body, into the dendritic arbor) and the retrograde direction (out of the arbor, back towards the cell body). At all ages, we measured balanced bidirectional transport (Figure 2E,G-H). Although anterograde mitochondria moved at significantly higher speeds than retrograde mitochondria (Figure 2F), there was no difference in linear flux rates (the number of motile mitochondria moving through the primary dendrite per minute) or mitochondrial lengths for anterograde versus retrograde mitochondria (Figure 2E,G). Thus, at any given age, equivalent volumes of mitochondria move in the anterograde and retrograde directions per unit time (Figure 2H). Comparing across ages, however, we measured a significant reduction in linear flux rates over time, with linear flux rates rapidly declining in early adulthood before plateauing in older flies (Figure 2E). In addition, we measured age-dependent changes in the size of the motile mitochondria, with mitochondrial lengths increasing with age up until 28 days before decreasing again in 56 day old flies (Figure 2G). This slight increase in mitochondrial size was not sufficient to compensate for the reduction in linear flux rates, however; overall, the total volume of mitochondria that exchanged through the primary dendrite declined with age (Figure 2H), with an approximately 5-fold reduction in 56 day old flies (volume exchange rate = ∼0.2 μm^3^/min) compared to 2 day old flies (∼1 μm^3^/min).

**Figure 2:**
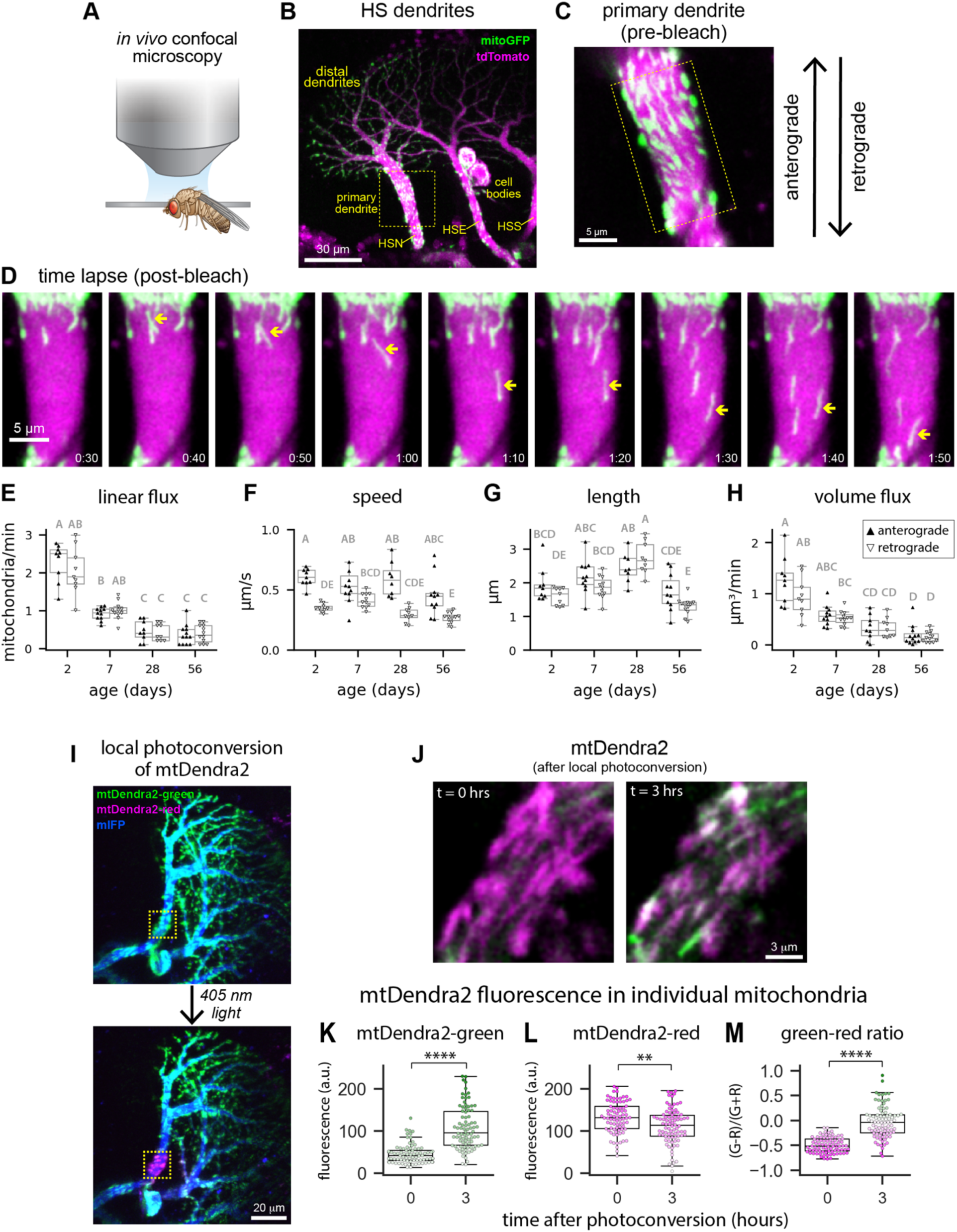
Mitochondrial dynamics in HS dendrites. A: Experimental setup. *In vivo* confocal imaging of head-fixed flies. B-C: Confocal images of mitochondria (mitoGFP, green) in HS dendrites (tdTomato, magenta) before photobleaching of mitochondria in the primary dendrite. The dashed yellow box in C indicates the primary dendrite shown in B. D: Image time series showing mitochondrial transport in the primary dendrite shown in C. To facilitate resolution of motile mitochondria, stationary mitochondria in the field of view were photobleached prior to image acquisition. The yellow arrow indicates a mitochondrion moving in the retrograde direction. Time (min:sec) is indicated in the bottom right corner of each image. E-H: Box plots showing the number of motile mitochondria moving through the primary dendrite per minute (linear flux, E), average pause-free speeds (F), lengths of motile mitochondria (G), and the total volume of mitochondria transported through the primary dendrite per minute (volume exchange, H), plotted as a function of fly age. Dots overlaid on the box plots indicate average values for individual flies (N = 9-11 flies per age). Letters indicate groups that are significantly different (Kruskal-Wallis test with post-hoc Dunn’s test, p < 0.05). I: *In vivo* confocal images of Dendra2-tagged mitochondria in HS dendrites before (top image) and after (bottom image) local photoconversion of mitochondria in the primary dendrite. J: Dendra2-tagged mitochondria in a primary HS dendrite immediately after photoconversion (left image) and three hours later (right image). K-M: Quantification of mtDendra2-green fluorescence (K), mtDendra2-red fluorescence (L), and a metric quantifying the green-to-red ratio (M) in individual mitochondria in primary HS dendrites after photoconversion. The green-to-red ratio is given by (G-R)/(G+R), where G is the green fluorescence intensity and R is the red fluorescence intensity. Dots overlaid on the box plots indicate measurements from individual mitochondria; dot color reflects the fluorescence intensity of each mitochondrion (K-L) or the ratio of green-to-red fluorescence within each mitochondrion (M). Asterisks indicate significant differences (Mann Whitney U test; ** p < 0.001, **** p < 10^-5^); N = 87 mitochondria (t = 0 hours) or 92 mitochondria (t = 3 hours) from six neurons. All box and whisker plots (in this figure and all subsequent figures) show the median, inter-quartile range (box), and 1.5 times the inter-quartile range (whiskers).

In the simplest mitostasis model, mitochondria carrying fresh proteins move in the anterograde direction, away from the soma, and replace old mitochondria in neurites which, in turn, are either degraded in place or trafficked back to the soma for degradation. Could the relatively low rates of mitochondrial transport in old flies impair the neuron’s ability to replenish mitochondria in distal dendrites? Based on our measurements of mitochondrial volume exchange through the primary dendrite and previously reported estimates of the total mitochondrial volume in the dendritic arbor (∼400 μm^3^) (Donovan et al., 2024), we estimated mitochondrial replacement timescales — i.e. the amount of time necessary to replace the entire volume of mitochondria in the dendrite, based on transport alone — as a function of fly age (see Methods). Even at reduced mitochondrial volume exchange rates in old flies (∼0.2 μm^3^/min, or ∼12 μm^3^/hr), we estimated a replacement timescale (∼1-2 days) that is substantially shorter than the median half-life for mitochondrial proteins in *Drosophila* (∼2 weeks) (Vincow et al., 2013). Thus, reduced mitochondrial transport rates in old flies are unlikely to limit the rate of mitochondrial replenishment in HS dendrites.

Having demonstrated that mitochondrial motility occurs at appreciable rates throughout adulthood, we next asked whether mitochondria in HS dendrites are also social, i.e. capable of fusing and dividing, as previously observed in neurons *ex vivo* (Fenton et al., 2024; Kochan et al., 2024; Lewis et al., 2018). To image mitochondrial fusion and fission events in HS dendrites, we combined *in vivo* confocal microscopy with HS-specific expression of a mitochondria-targeted photoconvertible fluorescent protein, mtDendra2. We converted a small population of Dendra2- tagged mitochondria from green to red fluorescence by locally illuminating a region of interest (ROI) in the dendrite with UV light, and then we imaged the photoconverted ROI over time (Figure 2I, S2A-B). We achieved efficient green-to-red photoconversion of the mtDendra2, and we were able to directly observe a small number of fusion events (Figure S2A-B). However, rapid photobleaching of mtDendra-green prevented us from imaging at a high frame rate for an extended period of time, hampering our ability to reliably image discrete fusion and fission events and quantify fusion/fission rates. We therefore measured mitochondrial content exchange in primary HS dendrites, rather than counting fusion/fission events, by quantifying green and red fluorescence within individual mitochondria in the photoconverted ROI immediately after photoconversion and three hours later (Figure 2J). We reasoned that if content exchange occurs in HS dendrites *in vivo*, then red mitochondria in the ROI should accumulate green fluorescence as mitochondria labeled with mtDendra2-green move into the ROI and fuse with stationary mitochondria labeled with mtDendra2-red. In contrast, if no exchange occurs and replenishment of mitochondria occurs via mitochondrial replacement alone — i.e. transport of new mitochondria into the dendrite balanced by transport of old mitochondria out of the dendrite — then green mitochondria should appear in the ROI and red mitochondria should disappear. We found, first, that mitochondria were primarily red immediately after photoconversion (Figure 2I-J,M). Three hours later, the amount of green-to-red fluorescence significantly increased in mitochondria due to an increase in mitoDendra2-green fluorescence as well as a slight decrease in mitoDendra2-red fluorescence (Figure 2K-M). Although a small number of mitochondria in the ROI three hours after photoconversion were almost entirely green, consistent with mitochondrial replacement, the majority were both green and red, consistent with mitochondrial content exchange (Figure 2J,M). We measured a similar pattern of green fluorescence recovery in HS distal dendrites, albeit at a slower rate than in primary dendrites (Figure S2C-F). These results demonstrate that recovery of mtDendra2-green in HS dendrites primarily occurs via fusion-mediated mitochondrial content exchange rather than replacement of whole mitochondria. Thus, mitochondria in HS dendrites are social as well as motile.

### Inhibition of mitochondrial transport and content exchange during development and adulthood disrupts dendrite form and function

Thus far, our results demonstrate that mitochondria in fully mature HS dendrites are dynamic: they exhibit bidirectional transport throughout the lifetime of the neuron, and they fuse and exchange their contents. Next, we wished to determine how deficits in mitochondrial transport or fusion, induced during development, affect mitochondrial localization, dendrite morphology, and dendrite function. Mitochondrial transport in neurons is mediated by an adaptor protein, Milton (TRAK1/2 in vertebrates), that links the mitochondrial outer membrane protein Miro to kinesin and dynein (Figure 3A) (Glater et al., 2006; Stowers et al., 2002). We knocked down expression of Milton in HS neurons during development and adulthood (using the R27B03 GAL4 driver, Figure S1) and measured the effects in young flies (7 days old). Milton knockdown abolished mitochondrial transport in HS primary dendrites, reducing linear flux in the anterograde and retrograde directions to zero (Figure 3B). Consistent with its effect on mitochondrial transport, Milton knockdown also caused mitochondrial mis-localization, with mitochondria enriched in the soma and depleted from the distal dendrites (Figure 3C-D). Milton knockdown also caused a significant reduction in HS dendrite size (i.e. reducing the total volume, length, and number of branches per dendrite) without affecting dendrite branch scaling (i.e. the relative widths of parent and daughter branches at each branch point) (Donovan et al., 2024) (Figure 3E-H, S3). Finally, Milton knockdown abolished calcium and ATP responses to preferred and null direction global motion (Figure 3I-J). Altogether, inhibition of mitochondrial transport, beginning prior to eclosion and continuing throughout adulthood, results in mitochondrial mis-localization and significant defects in HS dendrite form and function.

**Figure 3:**
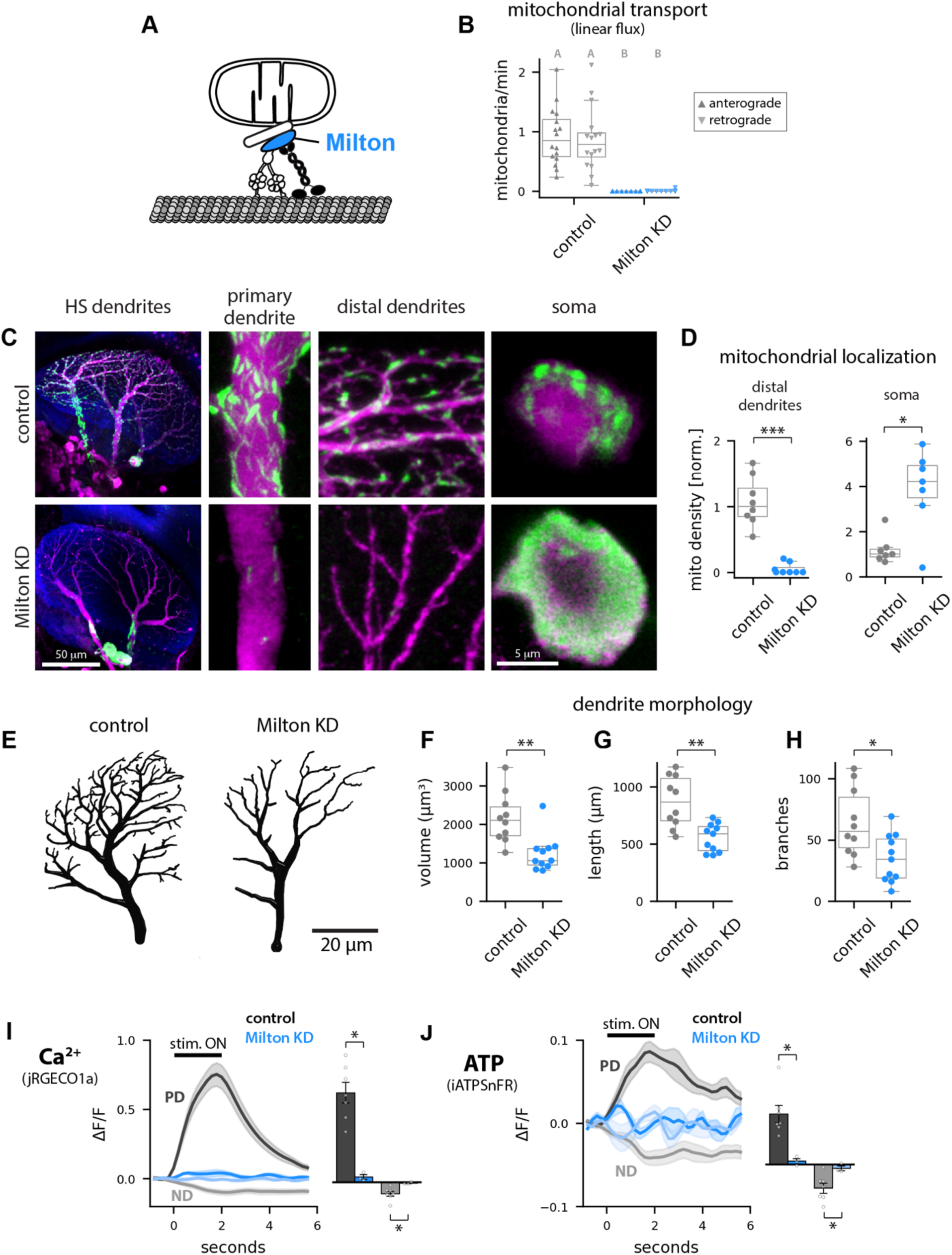
Milton knockdown during development and adulthood causes mitochondrial mis- localization and disrupts dendrite form and function. A: An adaptor protein, Milton (bright blue), links mitochondria to microtubule motor proteins via direct binding to a mitochondrial outer membrane protein (Miro). B: Mitochondrial linear flux rates in primary dendrites of control, and Milton knockdown samples. Dots overlaid on the box plots indicate measurements for individual cells; N = 16-17 cells per genotype. Letters indicate groups that are significantly different (Kruskal-Wallis test with post-hoc Dunn’s test, p < 10^-4^). C: Representative images of HS dendrites (labelled with cytosolic tdTomato, magenta) and the mitochondria within them (mitoGFP, green) in control (top row of images) and Milton knockdown samples (bottom row). Overview images on the left show all HS dendrites in the optic lobe; images of the primary and distal dendrites and somas are enlarged on the right. D: Mitochondrial volume densities, normalized to the median of the control, in distal dendrites (left plot) and somas (right plot). Dots overlaid on boxplots indicate average measurements for individual flies; N = 8 (distal dendrites) and 7 (somas) flies. Asterisks indicate significant differences (Mann Whitney U test; * p < 0.05, *** p < 0.001). E: Representative dendrite reconstructions from control and Milton knockdown neurons. F-H: Dendrite volume (F), dendrite length (G), and number of branches (H). Dots overlaid indicate measurements for individual dendrites; N = 10-11 dendrites per genotype. Asterisks indicate significant differences (Mann Whitney U test; * p < 0.05, ** p < 0.01). I-J: Average calcium (jRGECO1a, I) and ATP (iATPSnFR, J) responses to wide field square wave gratings moving in the preferred direction (PD) or null direction (ND) in distal dendrites of control (gray, N = 7 flies) and Milton knockdown (blue, N = 3 flies) samples. Insets show average response amplitudes. Asterisks indicate significant differences (Mann Whitney U test; * p < 0.05).

Next, to determine how disrupting mitochondrial content exchange during development and adulthood affects HS dendrites, we manipulated mitochondrial fusion by overexpressing or knocking down (via RNAi) the mitochondrial GTPase that mediates outer mitochondrial membrane (OMM) fusion, Marf (Mfn2 in vertebrates). Marf/Mfn2 drives OMM fusion via homotypic dimerization on adjacent mitochondria, and subsequent fusion of the inner mitochondrial membrane (IMM), mediated by the GTPase Opa1, results in exchange of mitochondrial matrix proteins and metabolites (Chan, 2020). Using local conversion of mtDendra2 in HS dendrites in young flies (7 days old), we found that Marf overexpression increased mitochondrial content exchange; ratios of green-to-red mitoDendra2 fluorescence within mitochondria in the photoconverted ROI were significantly higher and less variable than in control samples three hours after photoconversion (Figure 4A-D). In Marf knockdown samples, in contrast, photoconverted mitochondria remained entirely red three hours after photoconversion (Figure 4A-D), indicating that Marf is necessary for mitochondrial content exchange.

**Figure 4:**
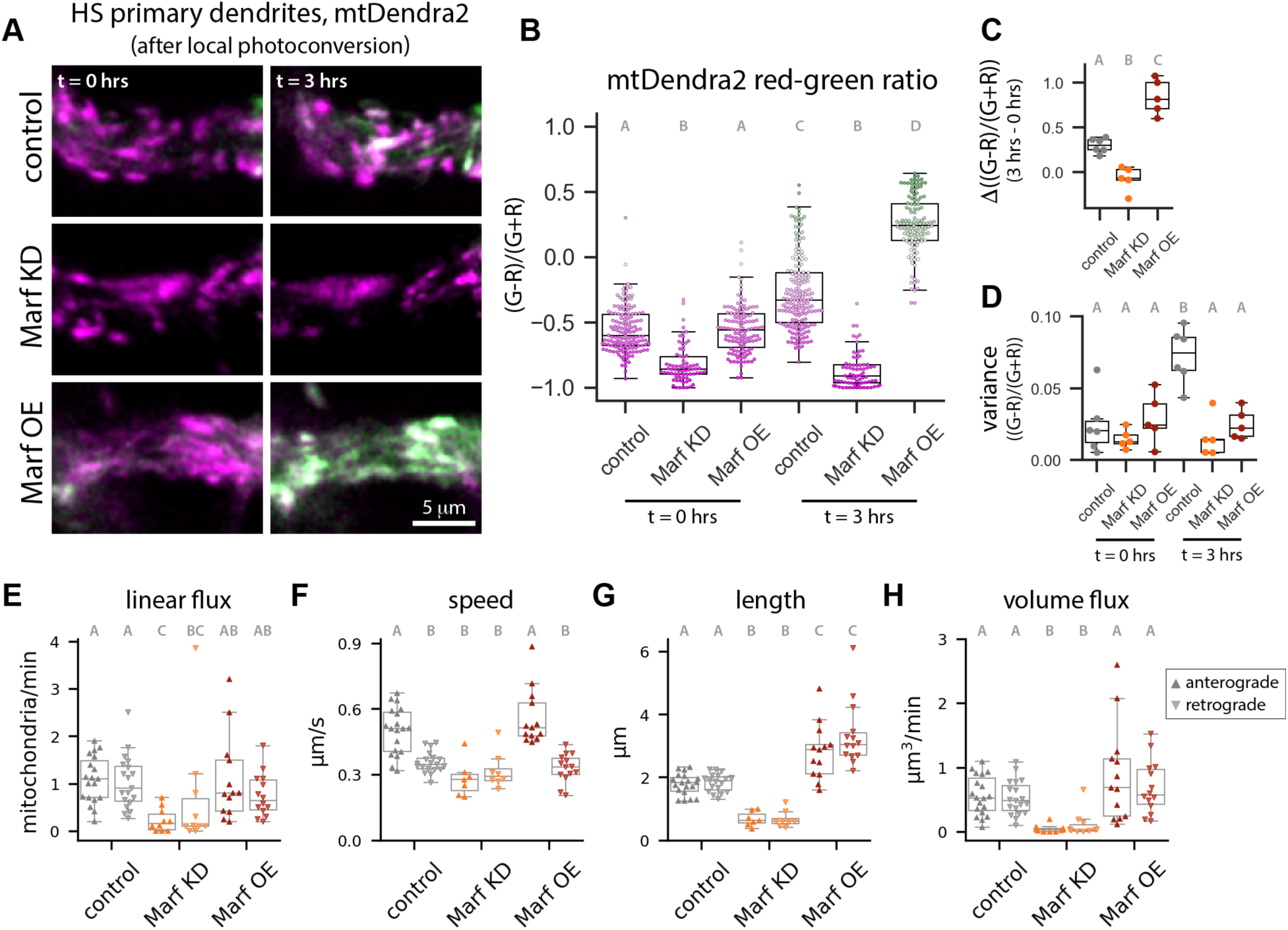
Marf is necessary for mitochondrial transport and content exchange. A: *In vivo* confocal images of Dendra2-tagged mitochondria in primary HS dendrites immediately after photoconversion (t = 0 hrs) and three hours later (t = 3 hrs) in control (top row), Marf KD (middle), and Marf OE (bottom) samples. B: Green-to-red ratio in individual mitochondria after photoconversion. Dots overlaid on the boxplots indicate measurements from individual mitochondria, and dot color reflects the green-to-red ratio within each mitochondrion. Letters indicate groups that are significantly different (Kruskal-Wallis test with post-hoc Dunn’s test, p < 10^-8^). C-D: Change in the average green-to-red ratio over time (3 hrs – 0 hrs after photoconversion) for the population of mitochondria within a primary HS dendrite (C), and the variation in the green-to-red ratio across the population at 0 and 3 hrs post photoconversion (D). Dots overlaid on the box plots indicate measurements from individual neurons; N = 5-6 neurons per genotype. Letters indicate groups that are significantly different (ANOVA with post-hoc Tukey’s test, p < 0.05). E-H: Quantification of mitochondrial motility. Box plots show linear flux rates (E), average pause-free speeds (F), mitochondrial lengths (G), and total volume of mitochondria transported through the primary dendrite per minute (H) for anterograde and retrograde transport; dots overlaid on the boxplots indicate average values for individual neurons (N = 10-19 neurons per genotype). Letters indicate groups that are significantly different (Kruskal-Wallis test with post-hoc Dunn’s test, p < 0.01).

In the absence of mitochondrial fusion, mitochondrial replenishment could in principle occur via transport-dependent mitochondrial replacement, with new mitochondria moving into the dendrite to replace old mitochondria. In our mtDendra2 experiments, we observed a small number of stationary green mitochondria in the photoconverted region three hours after photoconversion in control samples (Figure 2J, 4A-B). In Marf knockdown samples, however, no green mitochondria arrived in the photoconverted region (Figure 4A-B), suggesting that Marf plays a role in mitochondrial transport as well as mitochondrial fusion. Consistent with this idea, knockout of the vertebrate homolog of Marf (Mfn2) reduced axonal transport of mitochondria in cultured neurons (Misko et al., 2010). Mfn2 has also been shown to immunoprecipitate with the vertebrate homologs of Milton and Miro (Misko et al., 2010), suggesting that Mfn2 regulates mitochondrial transport via direct interaction with transport machinery. To measure the effects of Marf manipulations on mitochondrial transport in HS dendrites, we imaged GFP-tagged mitochondria moving through primary dendrites. On average, motile mitochondria in control samples were ∼2 μm long (Figure 2G, 4G). Marf knockdown significantly reduced the size of motile mitochondria (average length ∼0.7 μm), and Marf overexpression had the opposite effect (average length ∼3 μm) (Figure 4G). Marf overexpression had no effect on mitochondrial transport rates, but Marf knockdown significantly reduced the speed of mitochondria moving in the anterograde direction, as well as significantly reducing linear flux rates in both directions (Figure 4E-F). Together, the reduction in the size and number of motile mitochondria upon Marf knockdown dramatically reduced the total volume of mitochondria exchanged through the primary dendrite per minute (Figure 4H). These results indicate that Marf is necessary for mitochondrial transport as well as mitochondrial content exchange in HS dendrites.

Having demonstrated that Marf is necessary for mitochondrial transport and content exchange, we next measured the effects of Marf knockdown on mitochondrial localization and dendrite architecture. Consistent with Marf’s role in OMM fusion, Marf knockdown resulted in mitochondrial fragmentation and Marf overexpression caused mitochondrial elongation (Figure 5A-B). Similar to Milton knockdown, Marf knockdown also disrupted mitochondrial localization, reducing the mitochondrial volume density in the distal dendrites while increasing the mitochondrial density in the soma (Figure 5C). In addition, we noted that Marf knockdown resulted in accumulation of fragmented mitochondria in the soma stalk, a thin projection connecting the soma to the primary dendrite (as is typical of *Drosophila* neurons) (Figure 5A). Also similar to Milton knockdown, Marf knockdown reduced HS dendrite complexity, albeit in a more variable fashion, without affecting dendritic branch scaling (Figure 5D-F, S4). We measured bimodal distributions of dendrite volume, length, and number of branches; approximately half of the dendrites in our Marf knockdown samples were smaller than control dendrites, whereas the other half were indistinguishable from controls (Figure S4). This bimodal distribution of dendrite sizes suggests that variable Marf knockdown caused a threshold effect, in which stronger (or earlier) knockdown reduced dendrite size in 7 day old flies but weaker (or later) knockdown did not.

**Figure 5:**
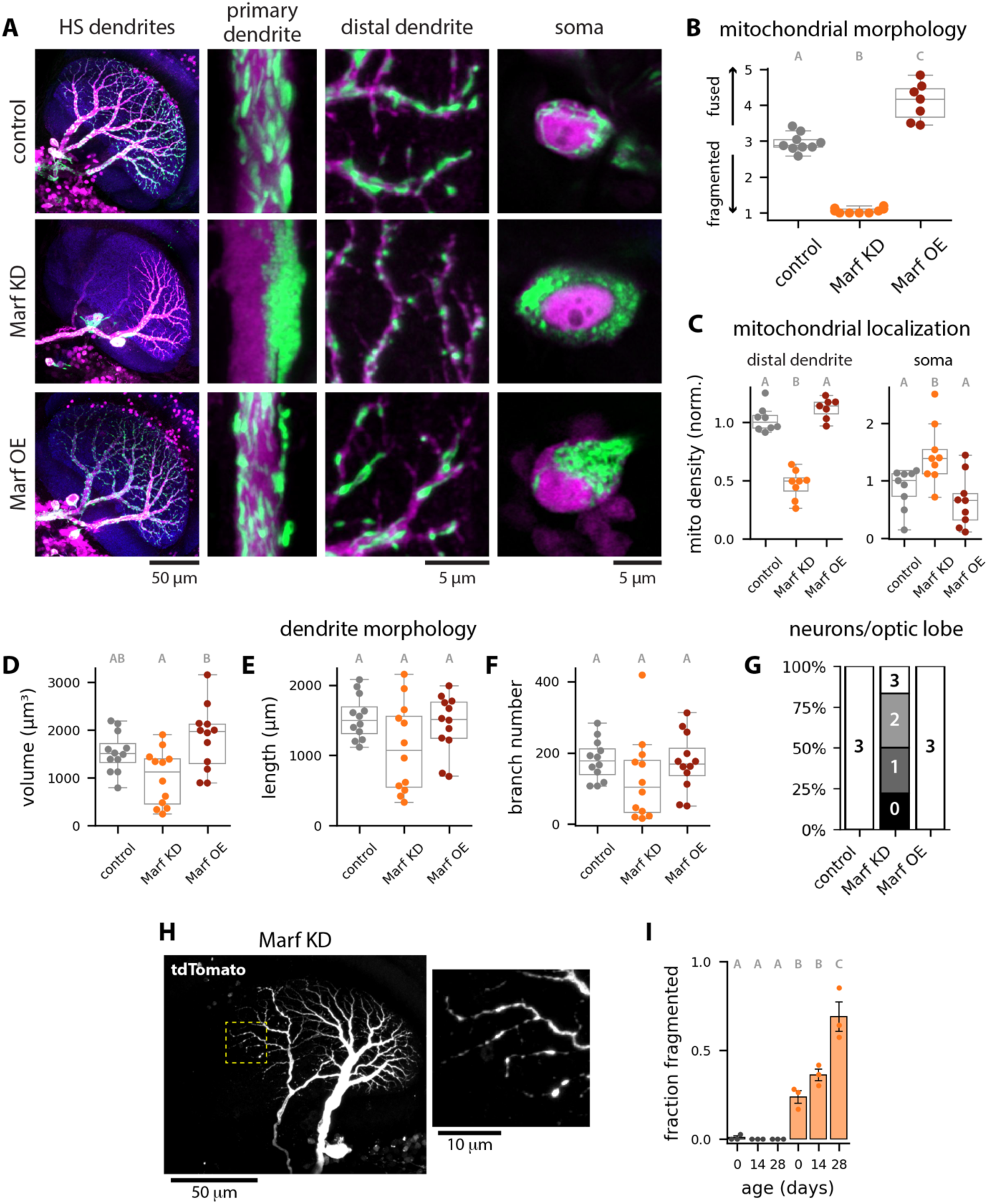
Marf knockdown during development and adulthood causes mitochondrial fragmentation and mis-localization, reduced cell numbers, and age-dependent dendrite degeneration. A: Representative images of HS dendrites (tdTomato, magenta) and the mitochondria within them (mitoGFP, green) in control (top row), Marf KD (middle) and Marf OE (bottom) samples. B: Mitochondrial morphology — fragmented versus fused — in HS distal dendrites. Dots overlaid on the box plots indicate measurements from individual flies (N = 7-9 flies per genotype). Letters indicate groups that are significantly different (Kruskal-Wallis test with post-hoc Dunn’s test, p < 0.05). C: Mitochondrial volume densities in distal dendrites (left) and somas (right). Dots overlaid on the box plots indicate average values for individual flies in the distal dendrites (N = 7-8 flies per genotype) or individual somas (N = 9 somas per genotype). Letters indicate groups that are significantly different (Kruskal-Wallis test with post-hoc Dunn’s test, p < 0.01). D-F: Dendrite volume (D), length (E), and total number of branches (F). Dots overlaid on the box plots indicate individual neurons (N = 12 neurons from 4-6 optic lobes). Letters indicate groups that are significantly different (Kruskal-Wallis test with post-hoc Dunn’s test, p < 0.05). G: Stacked bar plot showing the number of HS neurons per optic lobe in seven-day old flies; N = 14-19 optic lobes per genotype. H: Representative image of HS dendrites from a 14 day old Marf KD sample; yellow dashed box indicates the region enlarged in the image on the right. I: Fraction of HS dendrites that are degenerated, plotted as a function of age. Dots overlaid on the bar plot indicate the fraction fragmented per replicate (N = 3 replicates; 5 or more optic lobes per replicate). Letters indicate groups that are significantly different from each other (ANOVA with post-hoc Tukey’s test, p<0.05).

Finally, Marf knockdown also reduced the number of HS dendrites per optic lobe (Figure 5G). In control flies, we observed three HS neurons in all optic lobes (N = 19 optic lobes, st. dev. = 0). Marf knockdown reduced the average number of HS dendrites per optic lobe by half (1.4 +/- 1 neurons/optic lobe, N = 18 optic lobes) in 7 day old flies. The effects of Marf knockdown on neuron number were apparent prior to eclosion (Figure S5) and the number of cells did not significantly decrease with age (average cell count at day 28 = 1.1 +/- 0.7 neurons/optic lobe, N = 22 optic lobes), suggesting the reduction in cell count is due to developmental effects. This reduction in cell count is consistent with recent evidence that mitochondrial fusion and fission regulate neurogenesis (Dubal et al., 2022; Iwata et al., 2020). In addition to reduced cell counts, we also observed severe dendrite degeneration, characterized by fragmentation of the dendrite, in a subset of Marf knockdown samples (Figure 5H). The fraction of fragmented neurons increased over time, from ∼20% within 24 hours of eclosion to ∼70% at 28 days (Figure 5I). Thus, Marf knockdown causes a reduction in cell count prior to eclosion as well as dendrite degeneration that increases with age.

Altogether, these results indicate that deficits in mitochondrial dynamics, when induced prior to eclosion, disrupt dendrite morphology and function.

### Dendrite morphology and visual stimulus-driven ATP responses are robust to adult-onset deficits in mitostasis

Deficits in mitochondrial transport and fusion could reduce dendrite complexity in young adult flies due to developmental defects (e.g. reduced dendrite growth), loss of mitostasis in the adult and subsequent dendrite degeneration, or both. To determine whether adult-onset defects in mitochondrial dynamics disrupt dendrite maintenance, we used Gal80^ts^ to restricted Gal4-driven expression in HS to adulthood (Figure 6A). We reared crosses at 18C to suppress expression of Milton or Marf RNAi during development, and we induced knockdown during adulthood by shifting the temperature to 29C after eclosion. Using a pan-neuronal driver, we confirmed that adult-restricted expression of Milton and Marf RNAi resulted in significant reductions in Milton and Marf transcript levels in fly heads (Figure S6A). Seven days after the induction of Milton knockdown in HS, we measured near-complete inhibition of mitochondrial transport through primary dendrites (Figure 6B). Inhibition of mitochondrial transport had no significant effect on mitochondrial localization at day 7 (Figure S6B-C), but mitochondria were depleted from the dendrite and retained in the soma over time, resulting in a significant loss of mitochondrial volume in the dendrite at day 28 (Figure 6C-D). Similarly, Marf knockdown had no effect on mitochondrial morphology or localization in young flies (7 days old) but reduced mitochondrial size in old flies (Figure 6C,E, S6B,D-E). Marf knockdown also resulted in mitochondrial mis-localization over time, with mitochondria depleted from the dendrites and enriched in the soma at day 28 (Figure 6F). These results demonstrate that ongoing mitochondrial transport and fusion are necessary for mitostasis in adult HS dendrites.

**Figure 6:**
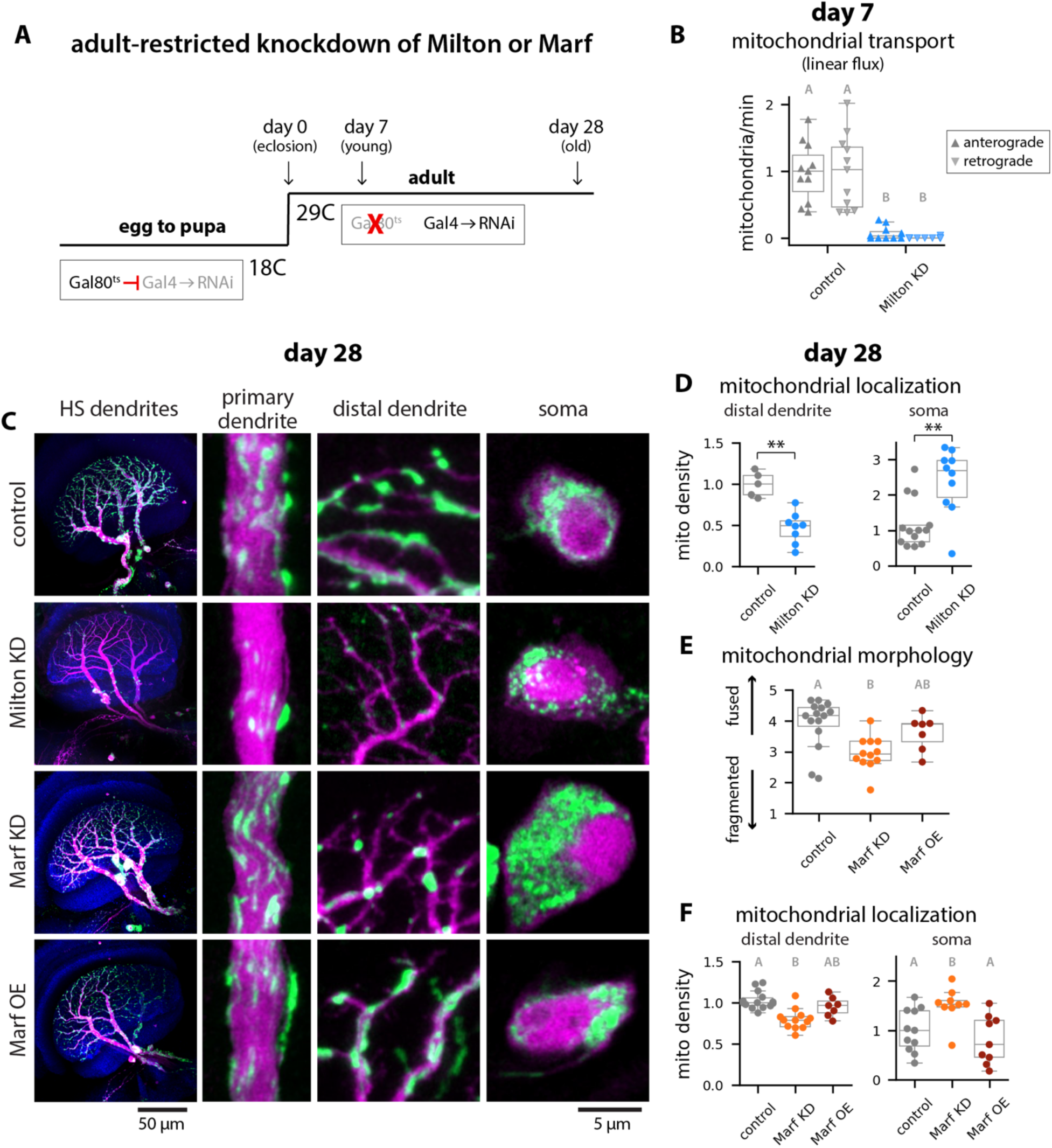
Adult-restricted knockdown of Milton or Marf depletes mitochondria from HS dendrites. A: Experimental approach. Gal80^ts^ suppressed Gal4-driven expression in crosses reared at 18C. After eclosion, adult flies were shifted to 29C to inactivate Gal80^ts^ and allow shRNAi expression in HS neurons. Experiments were conducted 7 or 28 days after the induction of shRNAi expression. B: Mitochondrial linear flux through primary HS dendrites in control and Milton KD samples at day 7. Dots overlaid on the box plots indicate measurements from individual flies (N = 11 flies per genotype). Letters indicate groups that are significantly different (Kruskal-Wallis test with post-hoc Dunn’s test, p < 0.001). C: Representative images of HS dendrites (tdTomato, magenta) and the mitochondria within them (mitoGFP, green) in control, Milton KD, Marf KD, and Marf OE samples at day 28. Overview images on the left show all HS dendrites in the optic lobe; images of the primary and distal dendrites and somas are enlarged on the right. D: Mitochondrial volume density in distal dendrites (left plot) and somas (right plot) at day 28 in control (gray) and Milton KD samples (blue). Dots overlaid on the box plots indicate average values for individual flies in the distal dendrites (N = 5-8 flies per genotype) or individual somas (N = 10-13 somas per genotype). Asterisks indicate significant differences (Mann Whitney U test, p < 0.01). E-F: Mitochondrial morphology — fragmented versus fused — in HS distal dendrites (E) and mitochondrial volume densities in distal dendrites and somas (F) at day 28 in control (gray), Marf KD (orange), and Marf OE (maroon) samples. Letters indicate groups that are significantly different (Kruskal-Wallis test with post-hoc Dunn’s test, p < 0.01).

However, despite causing mitochondrial mis-localization in old flies, neither Milton nor Marf adult-restricted knockdown affected the number or morphology of HS dendrites at the same time point (Figure 7A-D, G-J). We observed three HS neurons per optic lobe for all samples 28 days post-knockdown. When restricted to the adult, neither Milton nor Marf knockdown had an effect on dendrite volume, length, or branch number (Figure 7B-D, H-J). Thus, HS dendrite morphology is robust to Milton and Marf knockdown in the adult, suggesting the reduced dendrite complexity and age-dependent dendrite degeneration we observed upon knockdown prior to eclosion (Figure 3, 5) are due to deficits established during development.

**Figure 7:**
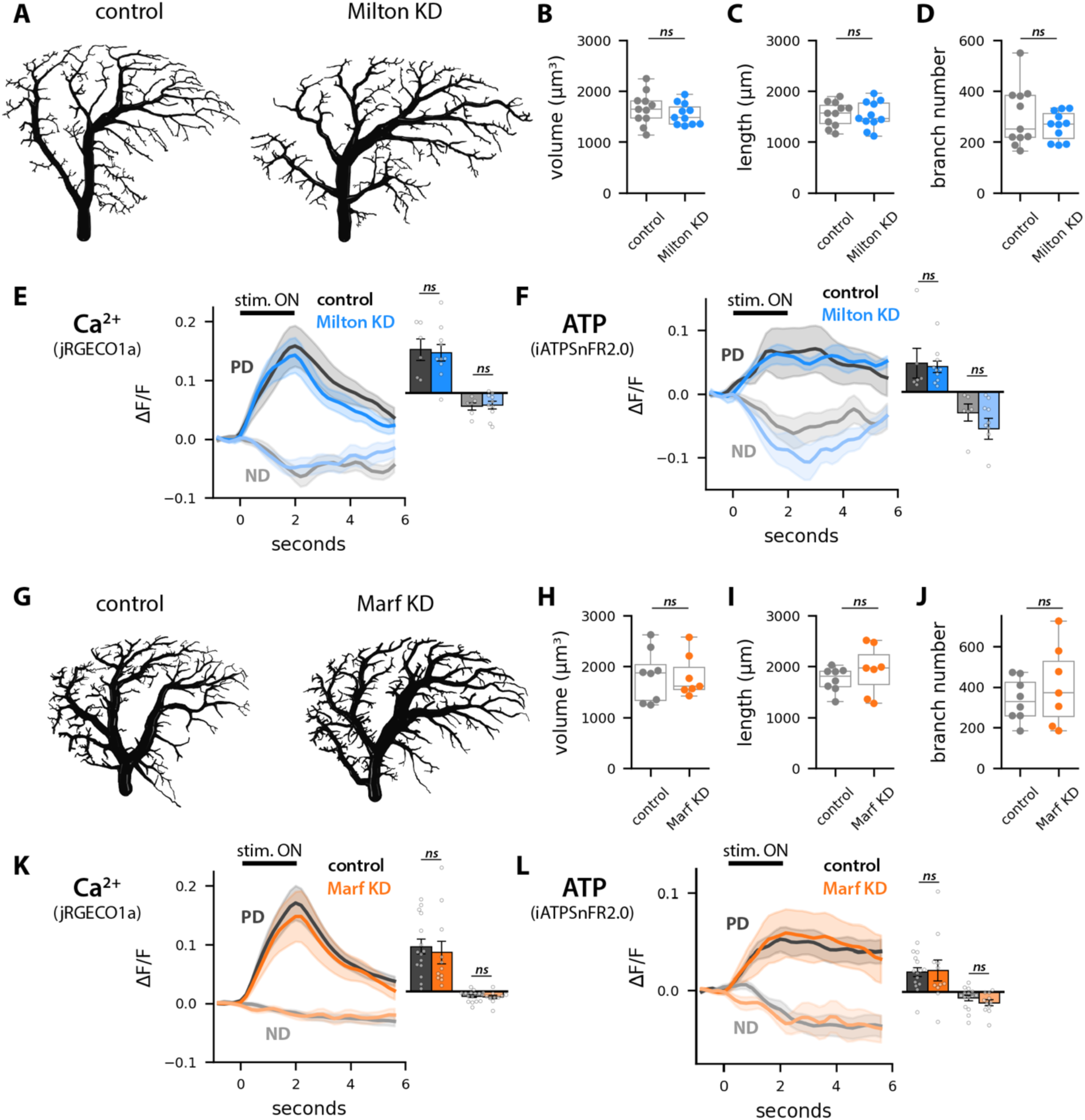
Adult-onset deficits in mitostasis do not affect HS dendrite morphology or visual stimulus-driven ATP responses. A: Representative reconstructions of control (left) and Milton knockdown (right) HS dendrites in 28 day old flies. B-D: Dendrite volume (B), length (C), and number of branch points (D) in control (gray) and Milton knockdown (blue) samples at day 28. Dots overlaid on the box plots indicate measurements for individual dendrites (N = 11 dendrites per genotype). E-F: Average calcium (jRGECO1a, E) and ATP (iATPSnFR2.0, F) responses to wide field square wave gratings moving in the preferred direction (PD) or null direction (ND) in distal dendrites of control (gray) and Milton knockdown (blue) samples at day 28. Insets show average response amplitudes. Shading on the line plots and error bars on the bar plots indicate the standard error of the mean; dots overlaid on the bar plots are average values for individual flies (N = 6-10 flies per genotype). G: Representative reconstructions of control (left) and Marf knockdown (right) HS dendrites in 28 day old flies. H-J: Dendrite volume (H), length (I), and number of branch points (J) in control (gray) and Marf knockdown (orange) samples at day 28 (N = 7-8 dendrites per genotype). K-L: Average calcium (jRGECO1a, K) and ATP (iATPSnFR2.0, L) responses to wide field square wave gratings moving in the preferred direction (PD) or null direction (ND) in distal dendrites of control (gray) and Marf knockdown (orange) samples (N = 11-15 flies per genotype). There are no significant differences between control and knockdown samples (Mann Whitney U test, p>0.05).

Finally, having found that HS dendrite morphology is unaffected by mitochondrial depletion during adulthood, we used *in vivo* two photon imaging of head-fixed flies to measure the effects of adult-restricted Milton or Marf knockdown on stimulus-driven calcium (jRGECO1a) and ATP (iATPSnFR2.0) responses. We presented full contrast moving square wave gratings on a screen positioned in front of the fly, with the grating moving either in the preferred direction (PD) or null direction (ND). Adult-restricted transport inhibition had no effect on mitochondrial localization in young flies (Figure S6B-C), and we were therefore unsurprised to find that Milton knockdown also had no effect on calcium or ATP responses at the same time point: PD motion drove increases in calcium and ATP levels, whereas ND motion had the opposite effect, in both control and Milton knockdown samples (Figure S6F-G). We were surprised, however, to find that although Milton knockdown depleted mitochondria from the dendrite in old flies (Figure 6C-D), this mitochondrial depletion had no effect on calcium or ATP responses (Figure 7E-F). In control dendrites, calcium response amplitudes declined with age (Figure 7E, S6F), and Milton knockdown neither exacerbated nor ameliorated this age-related decline: stimulus-driven calcium signals in 28 day old Milton knockdown samples were indistinguishable from controls (Figure 7E). Milton knockdown also had no effect on PD motion-driven increases in ATP levels (Figure 7F), suggesting that depletion of mitochondria does not negatively impact the dendrite’s ability to upregulate ATP production in response to excitatory input. In addition, although ND motion drove slightly larger reductions in ATP levels in Milton knockdown samples compared to controls, suggesting that mitochondrial depletion from the distal dendrite may cause ATP consumption to further outpace production during stimulation, this difference was not statistically significant (Figure 7F). Finally, adult-restricted Marf knockdown also had no measurable effect on stimulus-driven calcium or ATP responses in 28 day old flies (Figure 7K-L), despite causing mitochondrial mis-localization and fragmentation at the same time point (Figure 6C, E-F). Altogether, these results demonstrate that in HS dendrites, visual stimulus-driven fluctuations in calcium and ATP are largely impervious to adult-onset deficits in mitochondrial dynamics and localization.

## DISCUSSION

In this study, we found that *Drosophila* HS dendrites are resilient to adult-onset deficits in mitochondrial dynamics. Whereas disruption of mitochondrial transport and content exchange during development caused a range of significant phenotypes — mitochondrial fragmentation and mis-localization, loss of dendrite complexity, reduced cell counts, age-related dendrite degeneration, and complete abrogation of visual stimulus-driven calcium and ATP fluctuations —adult-restricted perturbations depleted mitochondria from the dendrite without affecting dendrite morphology or stimulus-driven calcium and ATP responses. Our results demonstrate that once properly developed, dendrites are capable of maintaining form and function *in vivo* in the absence of precisely positioned mitochondria.

We show here that mitochondria in fully developed HS dendrites are highly dynamic, exhibiting substantial bidirectional transport as well as fusion-mediated content exchange. Moreover, we demonstrate that ongoing mitochondrial transport is necessary for mitostasis: adult-restricted inhibition of mitochondrial transport (via Milton knockdown) caused gradual depletion of mitochondria from distal HS dendrites, with mitochondrial volume density reduced by ∼50% four weeks after induction of the knockdown. This timescale for mitochondrial depletion is roughly equivalent to the timescale for mitochondrial protein turnover in *Drosophila* (median mitochondrial protein half-life = ∼2 weeks) (Vincow et al., 2013), suggesting that absent mitochondrial transport, basal rates of mitochondrial degradation result in local loss of mitochondrial mass over time. Moreover, although some previously published measurements suggest that old, depolarized mitochondria are selectively trafficked back to the soma for degradation (Lin et al., 2017; Miller and Sheetz, 2004), adult-restricted Milton knockdown abolished all mitochondrial transport, including retrograde transport to the soma. Distal depletion of mitochondria is therefore likely due to local degradation of mitochondria, either via piecemeal degradation of mitochondrial proteins and mitochondria-derived vesicles and/or via local mitophagy or macroautophagy followed by retrograde trafficking of autophagosomes (Ashrafi et al., 2014; Cason et al., 2022; Goldsmith et al., 2022; Maday et al., 2012; Misgeld and Schwarz, 2017).

Despite causing mitochondrial depletion from the dendrite, adult-restricted inhibition of mitochondrial transport or fusion had no effect on HS dendrite morphology, indicating that precise mitochondrial positioning is not strictly necessary for neuronal maintenance *in vivo*. In mice, some neuronal cell types can also maintain normal morphologies for extended periods of time upon mitochondrial mis-localization (Berthet et al., 2014; Lopez-Domenech et al., 2016). In cortical pyramidal neurons, postnatal inhibition of mitochondrial transport (via knockout of Miro1) caused significant depletion of mitochondria from dendrites by four months of age, with a sizable fraction of dendritic branches entirely devoid of mitochondria (Lopez-Domenech et al., 2016). Despite this mitochondrial depletion, dendrite morphologies were indistinguishable from controls at four months. Morphological defects did eventually appear — reduced dendrite branching and neuronal degeneration occurred by 8 and 12 months of age, respectively — but the delayed onset of these defects, relative to the timing of mitochondrial depletion, indicates that pyramidal neurons are able to maintain normal dendrite morphology in the absence of mitochondria for a surprisingly long time (months). In addition to depending on age, the consequences of mitochondrial depletion also appear to depend on neuronal cell type. In dopaminergic (DA) neurons, postnatal knockout of the fission factor Drp1 depleted mitochondria from axons throughout the mouse midbrain but only caused degeneration in a subset of DA neurons, with susceptible neurons enriched in the substantia nigra and resilient neurons enriched in the ventral tegmental area (Berthet et al., 2014). The idea that susceptibility to mitochondrial perturbations varies with neuronal cell type is consistent with neurodegenerative disease phenotypes. Heterozygous mutations in different fusion factors affect distinct populations of neurons: Mfn2 mutations are associated with axonopathy of peripheral motor and sensory neurons in Charcot-Marie-Tooth disease type 2A (Zuchner et al., 2004), whereas Opa1 mutations are associated with progressive degeneration of retinal ganglion cells in dominant optic atrophy (Delettre et al., 2000). Similarly, mutations in master regulators of mitophagy, Pink1 and Parkin, cause selective degeneration of DA neurons in the substantia nigra in Parkinson’s disease (Pickrell and Youle, 2015). It remains unclear, however, why these mitochondrial mutations perturb specific subsets of neurons while other neuronal cell types are unaffected.

Neuronal susceptibility versus resilience to mitochondrial insults may be due to cell type-dependent differences in energetic demands (Alle et al., 2009; Carter and Bean, 2009; Niven, 2016). In addition, variation in alternative power management mechanisms (e.g. dynamic regulation of glycolysis rather than OXPHOS) could confer resilience to mitochondrial depletion or dysfunction in a cell type-specific fashion. In HS dendrites, adult-onset mitochondrial depletion had no significant effect on visual stimulus-driven increases in ATP, suggesting that HS dendrites do not rely on mitochondrial OXPHOS for local, acute regulation of ATP production. Instead, visual stimulus-driven fluctuations in ATP levels may be due to activity-dependent regulation of glycolytic flux. Consistent with this idea, blocking glycolysis causes a significant reduction in ATP levels upon neuronal activation in culture (Rangaraju et al., 2014). Moreover, recent work has shown that neurons in *C. elegans* tune glycolysis rates in a cell type-dependent fashion (Wolfe et al., 2024), as well as localizing glycolytic enzymes to synapses (Jang et al., 2016) and upregulating glycolytic flux under hypoxic conditions (Wolfe et al., 2024). In circadian clock neurons in *Drosophila*, cell type-specific knockout of either Marf or Opa1 triggers increased expression of glycolysis enzymes (Richhariya et al., 2025), suggesting that neurons can compensate for mitochondrial dysfunction via upregulation of glycolysis. Thus, although oxidative phosphorylation is the most efficient mechanism for converting high-energy carbon compounds into ATP, dynamic regulation of glycolysis may be sufficient to meet local, acute increases in ATP demand upon neuronal activation.

HS dendrites may also rely on spatiotemporal energy buffering, mediated by metabolically inert phosphagens, to ensure sufficient global supply of ATP throughout the neuron even when mitochondria are mis-localized. In phosphagen buffering systems, which have been extensively studied in muscle, creatine kinase (or arginine kinase, in flies) is functionally coupled with glycolysis or oxidative phosphorylation (Wallimann et al., 1992). At sites of high ATP production, creatine kinase catalyzes the transfer of the terminal phosphate group of ATP to creatine (or arginine, in flies), followed by diffusion of phosphocreatine throughout the cell. The resultant global pool of phosphocreatine, maintained in excess of ATP, allows creatine kinase to efficiently transfer the phosphate back to ADP at sites of high ATP consumption, thereby buffering fluctuations in ATP in space and time. In our experiments, inhibition of mitochondrial transport results in redistribution of mitochondria, with mitochondria enriched in the soma and depleted from the dendrite. This spatial distribution of mitochondria is reminiscent of mitochondrial localization in photoreceptors in salamanders and zebrafish, in which mitochondria are enriched in photoreceptor inner segments and excluded from synaptic terminals (Linton et al., 2010). In these photoreceptors, energy (in the form of phosphocreatine) flows from mitochondria toward synaptic terminals. In HS neurons, energetic flux from the soma to the distal dendrites, mediated by the *Drosophila* phosphagen system (arginine kinase/phosphoarginine), may protect the dendrite from degeneration when mitochondria are mis-localized.

Altogether, our work suggests that dynamic regulation of ATP production by precisely positioned mitochondria is not strictly necessary to match energy supply with demand and maintain neuronal form and function, highlighting the need to define alternative power management mechanisms in neurons *in vivo*.

## ACKNOWLEDGEMENTS

We thank Catherine Collins for providing UAS-mtDendra2 flies and Tom Clandinin for providing UAS-iATPSnFR2.0 flies. We are grateful to Franck Polleux, Mimi Shirasu-Hiza, and all members of the Barnhart lab for comments on the manuscript and helpful discussions. This work was supported by the NIH (R01NS121179 to ELB and F31NS129199 to EDB) and the NSF (grant number 2227609 to ELB).

## AUTHOR CONTRIBUTIONS

Conceptualization, ELB, HQW, DGF, and EDB; Investigation, HQW, DGF, EDB, HSW, BMR, and LB; Software AJA, JIK, GN, and ELB; Formal Analysis, HQW, DGF, EDB, HSB, and ELB; Writing — Original Draft, HQW, DGF, and EBD; Writing — Review and Editing; ELB, HQW, DGF, and EDB; Supervision, ELB.

## DECLARATION OF INTERESTS

The authors declare no competing interests.

## METHODS

| REAGENT or RESOURCE | SOURCE | IDENTIFIER |
| --- | --- | --- |
| **Antibodies** |  |  |
| Chicken polyclonal anti-GFP | Aves Labs | GFP-1010; RRID: AB_2307313 |
| Rabbit polyclonal anti-RFP | Rockland | 600-401-379; RRID: AB_2209751 |
| Mouse monoclonal anti-BRP | DSHB | nc-82; RRID: AB_2314866 |
| Alexa Fluor 488 Goat anti-Chicken IgY | Thermo Fisher Scientific | A-11039; RRID: AB_2534096 |
| Alexa Fluor 555 Goat anti-Rabbit IgG | Thermo Fisher Scientific | A-21428; RRID: AB_2535849 |
| Alexa Fluor 647 Goat anti-Mouse IgG | Thermo Fisher Scientific | A-21235; RRID: AB_2535804 |
| **Chemicals** |  |  |
| Bondic UV glue | BONDIC | N/A |
| ProLong™ Gold Antifade Mountant | Invitrogen | S36937 |
| Paraformaldehyde 16% | Electron Microscopy Sciences | 15710 |
| Triton X-100 | Biomatik | A4025 |
| TRIGent (TRIzol) | Biomatik | K5161 |
| Chloroform | Sigma-Aldrich | 288306 |
| Ultrapure DNase/RNase Free Distilled Water | Thermo Fisher Scientific | 10-977-015 |
| 2-Propanol | Sigma-Aldrich | 190764 |
| Ethanol | EMD Millipore | 108543 |
| RevertAid First Strand cDNA Synthesis Kit | Thermo Fisher Scientific | K1621 |
| Invitrogen™ DNase I, Amplification Grade | Invitrogen | 18-068-015 |
| SYBR™ Green Universal Master Mix | Thermo Fisher Scientific | 4309155 |
| Custom 25 nmole DNA Oligos (qRT-PCR Primers) | IDT Technologies | N/A |
| **Experimental models: organisms/strains** |  |  |
| Canton-S | Bloomington *Drosophila* Stock Center (BDSC) | 64349 |
| GMR27B03-GAL4 (HS) | BDSC | 49211 |
| GMR57C10-GAL4 (nSyb) | BDSC | 39171 |
| w+; UAS-mitoGFP,UAS-tdTomato/CyO; GMR27B03-GAL4/TM6B | Donovan et al. | N/A |
| UAS-mtDendra2 | Gift from Catherine Collins, Case Western University | N/A |
| UAS-mIFP-T2A-HO1 | BDSC | 64181 |
| w+: UAS-mtDendra2/CyO; GMR27B03-GAL4, UAS-mIFP-T2A-HO1/TM6B | This study | N/A |
| UAS-jRGECO1a/TM6B | BDSC | 63794 |
| UAS-iATPSnFR2.0 | Gift from Tom Clandinin, Stanford University | N/A |
| w+; UAS-iATPSnFR2.0/CyO; GMR27B03-GAL4, UAS-jRGECO1a/TM6B | This study | N/A |
| UAS-Milt RNAi | BDSC | 44477 |
| UAS-Marf.RNAi | BDSC | 67158 |
| UAS-Marf.HA/Cyo:TM6B | BDSC | 67157 |
| tubP-GAL80<sup>ts</sup>; TM2/TM6B | BDSC | 7108 |
| **Software and algorithms** |  |  |
| Fiji | Schindelin et al. | fiji.sc |
| ilastik | Berg et al. | www.ilastik.org |
| MATLAB | MathWorks | www.mathworks.com |
| PsychoPy | Peirce et al. | www.psychopy.org |
| Python3 | Python | www.anaconda.com/products/ distribution |
| neuTube | Feng et al. | neutracing.com |
| nnUNET | Isensee et al. | github.com/MIC-DKFZ/nnUNet |
| Custom written Python code for quantifying mitochondrial transport, mitochondrial mixing, mitochondrial morphology, dendrite fragmentation, and extracting visual stimulus-driven calcium signals from *in vivo* images | this paper |  |
| Custom written MATLAB code for quantifying neuronal morphology | Donovan et al. |  |

### *Drosophila* strains and husbandry

Flies were reared on standard molasses cornmeal under a regular 12h light/dark cycle. Unless indicated otherwise, crosses were reared at 25°C. To restrict expression of transgenes to the adult using Gal80^ts^, crosses were reared at 18°C, and progeny were collected at the day of eclosion and moved to 29°C. All crosses were flipped onto fresh food every 3-4 days. All *Drosophila* strains used in this work are listed in the table above.

### *Drosophila* whole brain dissection and immunostaining

Whole brains were dissected from pupae and adult female flies in cold 2% paraformaldehyde supplemented with 0.1M L-Lysine, followed by fixation for 1 hour on ice; pupal brains were briefly (∼1 minute) rinsed in PBS with 1% Triton prior to fixation. After fixation, all brains (adult and pupal) were washed using PBST (PBS with 0.5% Triton) three times at 1 minute intervals and blocked in PBST-NGS (PBST with 5% normal goat serum, Abcam) for 30 minutes at room temperature. Brains were then incubated in primary antibodies diluted in PBST-NGS (Chicken anti-GFP, 1:1000 dilution; Rabbit anti-RFP, 1:50; and Mouse anti-BRP, 1:10) for 48 hours at 4°C. Brains were then washed three times at 30 minute intervals with PBST-NGS before incubation with secondary antibodies (Alexa-488 Goat anti-Chicken, 1:1000; Alexa-555 Goat anti-Rabbit, 1:500; and Alexa-647 Goat anti-Mouse, 1:500) for 24 hours at 4°C. After staining, brains were washed with PBST-NGS three times at 20 minute intervals and mounted on microscope slides. Each brain was positioned in 15uL of SlowFade Gold Antifade (Invitrogen) in an iSpacer (Sun Jin Lab) with the posterior side of the optic lobes oriented towards the coverslip.

Immunostained brains were imaged using a either a Leica SP8 confocal microscope (images in Figure 3) equipped with a 63X 1.4 NA oil immersion objective (Leica) or a confocal Zeiss LSM900 (all other images) equipped with an Airyscan detector (Zeiss) and an 63X 1.4 NA oil immersion objective (Zeiss). For samples imaged with the Leica, overview z-stacks of the entire lobula plate (voxel size: 137nm x 137nm x 500nm) were acquired for each sample, along with higher resolution z-stacks of somas and primary and distal dendrites (voxel size: 49.1nm x 49.1nm x 500nm). On the Zeiss, overview z-stacks (voxel size: 49.5nm x 49.5nm x 500nm) and higher resolution z-stacks of the subcellular compartments (voxel size: 42.7nm x 42.7nm x 150nm) were collected using Airyscan Multiplex mode with 2x line averaging.

### In vivo imaging

Female flies were anesthetized on ice and positioned in a keyhole cut in a thin metal shim. Flies were positioned such that the back of the head was flush with the shim and the eyes were below the shim. Flies were then secured with UV-cured glue (Bondic) and dissected in a cold sugar saline solution (103 mM NaCl, 3 mM KCl, 5 mM TES, 1 mM NaH2PO4, 26 mM NaHCO2, 4 mM MgCl2, 1.5 mM CaCl2, 10 mM trehalose, 10 mM glucose, and 7 mM sucrose, bubbled with carbogen to reach a pH of 7.2). Fine forceps were used to create a small opening in the cuticle and remove fat and trachea above the left optic lobe. Brains were then perfused with room-temperature saline and imaged using an integrated confocal and two-photon microscope (Leica SP8 CSU MP Dual) equipped with a 20X 1.0 NA water immersion objective (Leica) and a resonant scan head.

For confocal imaging of mitochondrial transport, GFP targeted to the mitochondrial matrix (mitoGFP) and a cytoplasmic red fluorescent protein (tdTomato) were selectively expressed in HS neurons using the R27B03-GAL4 driver line. Stationary mitochondria within the primary HS dendrite were photobleached to facilitate resolution of individual motile mitochondria, and confocal z-stacks (voxel size = 108.5nm x 108.5nm x 1μm; field of view = 26.1μm x 26.1μm x ∼13μm) were collected at five second intervals for five to ten minutes.

For confocal imaging of mitochondrial mixing, a photoconvertible fluorescent protein targeted to mitochondria (mitoDendra2) and a far-red fluorescent protein expressed in the cytosol (mIFP) were selectively expressed in HS neurons. Dendra2-labeled mitochondria within a 10.4μm x 10.4μm region of interest (ROI) were converted from green to red fluorescence by illumination with a 405nm laser for ∼10 seconds. Confocal z-stacks (voxel size = 108.5nm x 108.5nm x 1μm; field of view = 56.2μm x 56.2μm x ∼30μm) centered on the ROI were imaged immediately before and after photoconversion as well as three hours after photoconversion. Samples were kept in the dark in a 25C incubator between time points. To observe individual fusion events (Figure S2), time lapse images of photoconverted ROIs were collected at 10 minute intervals for 90 minutes.

For two-photon imaging, a green fluorescent ATP reporter (iATPSnFR2.0) and a red fluorescent calcium indicator (jRGECO1a) were expressed in HS neurons and simultaneously excited using a tunable MP laser set to 920nm (to excite iATPSnFR) and a fixed 1045nm laser line (to excited jRGECO1a) (Spectra-Physics Insight X3 Dual). Images (pixel size: 124nm x 124nm; field of view: ∼25μm x 12μm) were collected at 100ms time intervals for 2 minutes. Multiple (∼2-5) distal dendrites were imaged per fly; total imaging time per fly was approximately 30 minutes.

### Visual stimulus presentation

Visual stimuli were created using PsychoPy and displayed via a digital light projector (DLP LightCrafter, Texas Instruments). The stimuli were projected onto a screen positioned ∼2 cm away from the fly’s eye, covering ∼60° of its visual field horizontally and ∼60° vertically. To prevent the microscope from detecting light emitted by the projector, a 472/30 nm bandpass filter (Semrock) was used to filter the stimulus before it was projected onto the screen. The stimulus was refreshed at 60 Hz. To synchronize the stimuli and the imaging frames, voltage signals from the microscope were relayed to PsychoPy through a LabJack device. To measure stimulus-evoked calcium and ATP responses in HS dendrites, full contrast widefield square wave gratings (λ = 30°) that spanned the entire stimulus screen and moved at 30°/s were presented for 2 seconds, with static gratings presented for 2 seconds before and after motion.

### Image analysis

#### Rapid classification of mitochondrial morphology and dendrite degeneration

To quantify mitochondrial morphology, images of mitochondria (labelled with mitoGFP) in HS distal dendrites were scored by eye on a five point scale ranging from highly fissed (1) to highly fused (5). To quantify dendrite degeneration, the fraction of neurons exhibiting severe dendrite fragmentation was scored based on images of HS distal dendrites (labelled with a cytosolic volume marker, tdTomato) for three independent replicates consisting of at least five optic lobes. In both cases, custom-written Python code was used facilitate scoring of randomly-presented images in a gamified fashion. Players first worked through a tutorial that defined image classes (e.g. fissed versus fused mitochondria and fragmented versus unfragmented dendrites) and indicated which keys to press for each image class. Then, players had to pass a graded trial run in which their scores for a subset of images were compared to gold standard scores (generated by the researcher who generated the images) before scoring all images. Images were stripped of identifying information (i.e. genotype) and presented in a randomized order. Each image was classified by at least four players, and final scores for each image were calculated by averaging the scores of all players.

#### Quantification of mitochondrial density

Mitochondrial density measurements were taken from 1- 3 z-stacks per compartment (distal dendrites or somas) for each optic lobe. Automatic semantic segmentation of mitochondria, neuron, and background pixels was carried out using ilastik (Berg et al., 2019) and nnUNET (Isensee et al., 2021). To generate training data, images were segmented in ilastik using pixel classification mode, allowing for labels of neuron, mitochondria, and background to be assigned to individual pixels. The segmented masks were then manually cleaned in ImageJ and used, along with the original images, to train an nnUNET model. The trained nnUNET model was then used to generate predictions for a larger set of images. When necessary, more masks were cleaned and fed into nnUNET for additional rounds of training. Following successful prediction, segmentations were manually cleaned to remove background signals. Finally, mitochondrial density measurements — the total number of mitochondrial pixels divided by the sum of the mitochondrial plus neuron pixels — were extracted segmented images using custom-written Python code.

#### Quantification of dendrite morphology

The number of HS dendrites was counted by manual scoring the number of primary dendrites per optic lobe. To measure dendrite morphology, three-dimension reconstructions of dendrites were generated based on a cytosolic volume marker (tdTomato) in a semi-manual fashion in neuTube (Feng et al., 2015). Before analysis, primary dendrite lengths were set to the length of the shortest primary dendrite across all genotypes and ages using TreesToolbox (Cuntz et al., 2011) in Matlab. Total dendrite volume, total dendrite length, number of branch points, parent-daughter and sister-sister scaling parameters were analyzed using previously published MATLAB and Python code (Donovan et al., 2024). To determine whether dendrite morphologies followed normal or bimodal distributions (Figure S4), measurements of dendrite volume, length, and branch numbers were fit to Gaussian mixture models using the Mclust function in R.

#### Quantification of mitochondrial transport/motility

Motile mitochondrial lengths, speeds, and linear flux rates were measured from maximum projections of confocal z-stack images of mitoGFP and cytosolic tdTomato, expressed in HS neurons. Prior to taking measurements, max projections of z-stack images were aligned using either TurboReg Fiji plugin or custom-built Python code. Mitochondrial lengths were measured using the line tool in Fiji. Individual mitochondria were hand-tracked for both the anterograde and retrograde direction using the Tracking Fiji built in plugin. Measurements of pause-free mitochondrial speeds and linear flux rates were extracted from mitochondrial trajectories using custom-written Python code. Mitochondrial speeds were defined as the average speed (distance over time) above an instantaneous speed threshold of 0.1 μm/s. Linear flux rates were defined as the number of motile mitochondria move through the primary dendrite in either the anterograde or retrograde direction per unit time. Volume fluxes (!_!_), or total mitochondrial volumes moving through the primary dendrite per unit time, were calculated based on measurements of linear flux rates (!_"_) and lengths of motile mitochondria (#) as well as a fixed mitochondrial radius ($ = 0.3 μm, estimated from previous measurements of mitochondria in electron microscopy images of HS primary dendrites (Donovan et al., 2024)): !_!_ = !_"_ × ($^#^#. The time scale) for mitochondrial replacement (i.e. the time required to replace the entire volume of mitochondria in an HS dendritic arbor via anterograde and retrograde transport alone) was estimated as a function of fly age based on previous measurements of the total mitochondrial volume in an HS dendrite (*_$%$_ = ∼400μm^3^) and average volume flux measurements:) = *_$%$_ ⁄!_!_ .

#### Quantification of mitochondrial content exchange

*In vivo* images of mtDendra2 were analyzed to measure mitochondrial content exchange in primary and distal HS dendrites. Confocal z slices collected at two points after local photoconversion of mtDendra2-green to mtDendra2-red (immediately after photoconversion (t0) and three hours later (t3)) were mutually aligned using custom-written Python code, and individual mitochondria in each z slice (2-3 slices per sample) were manually segmented using the paint tool in Fiji. Then, using custom-written Python code, average green and red fluorescence intensities were extracted from each mitochondrion, and the ratio of green (G) to red (R) fluorescence was calculated as (G-R)/(G+R). The change in this green-red ratio over time was calculated for photoconverted ROIs in each primary dendrite by subtracting the average green-red ratio for all mitochondria in the ROI at t0 from the average green-red ratio at t3. At each time point, variance in the green-red ratio was calculated according to 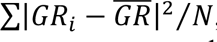, where ./_&_ is the green-red ratio for an individual mitochondrion, .*GR* is the average green-red ratio, and 2 is the number of mitochondria in the photoconverted ROI.

#### Quantification of visual stimulus-driven calcium and ATP responses

To measure calcium (jRGECO1a) and ATP (iATPSnFR2) signals in HS dendrites, time-series images were aligned using Python code based on the ImageJ TurboReg plugin. Then, ROIs were masked from the average projection of the time series via simple thresholding and manual clean-up in Fiji. Custom-written Python code was then used to measure pixel intensities within each ROI. To measure average responses moving square wave gratings, images were binned first by direction of motion (PD versus ND) and then by time (relative to the onset of motion). ΔF/F was calculated as (F(t)- F0)/F0, where F0 was the average fluorescence intensity in the one second immediately prior to the onset of motion. Response amplitudes were calculated by summing ΔF/F values in a three second time window immediately following the onset of motion; this time window includes the two second stimulus period as well as the second immediately following the cessation of motion. The direction selectivity index (DSI) was calculated according to DSI = ½ (Rpref-Rnull)/Rpref, where Rpref is the amplitude of the response to the preferred direction of motion and Rnull is the amplitude of the response to motion in the opposite direction.

#### RNAi validation

To validate the adult-restricted, RNAi-mediated knockdown of Milton or Marf, shRNAi were pan-neuronally expressed using the nSyb-GAL4 driver (GMR57C10-GAL4). Expression was suppressed during development using Gal80ts. Crosses were reared at 18C, and adult flies were shifted to 29C within 24 hours of eclosion. Transcipt levels were measured using qRT-PCR seven days after the induction of the knockdown. For each biological replicate, 25 fly heads were separated from the body and homogenized in 500 μl of TRIzol reagent (Biomatik) for RNA extraction, followed by the addition of 100 μl chloroform and subsequent extraction steps per manufacturer protocol. Samples were treated with DNase for RNA purification and cDNA was synthesized using the Revertaid First Strand cDNA Synthesis Kit (Thermo Scientific). Quantitative RT-PCR was performed on the StepOnePlus Applied Biosystems Real Time PCR System using the Applied Biosystems SYBR Green PCR Master Mix (Thermo Scientific). The transcript levels were normalized to the reference gene (Actin5C) and quantified relative to the control using the delta-delta Ct method, providing relative fold change.

The following primer pairs (IDT Technologies) were used: Actin5c-fwd- TTGTCTGGGCAAGAGGATCAG Actin5c-rev- ACCACTCGCACTTGCACTTTC Milton-fwd- GCAGACGATGGCACAGATACT Milton-rev- CGTCGAGCAGGGAGTTGAC Marf-fwd-ACTCATCGCTGCAACAATCC Marf-rev-ATCTGGAGCGGTGATTTGTCG

### Statistical analysis

All datasets were tested for normality using the Shapiro-Wilk test. For normally-distributed data, single comparisons were performed using paired or unpaired T tests; multiple comparisons were performed using ANOVA and post-hoc Tukey’s tests. For non-normal data, single comparisons were performed using Mann Whitney U tests; multiple comparisons were performed using the Kruskal-Wallis test, followed by Dunn’s test with Benjamini-Hochberg p value adjustment. The specific tests used for each dataset are indicated in figure legends. All statistical tests were implemented using the stats module in SciPy.

## Notes

### Competing Interest Statement

The authors have declared no competing interest.

