## Supplemental Figures for "*Drosophila* HS dendrites are resilient to adult-onset deficits in mitochondrial dynamics"

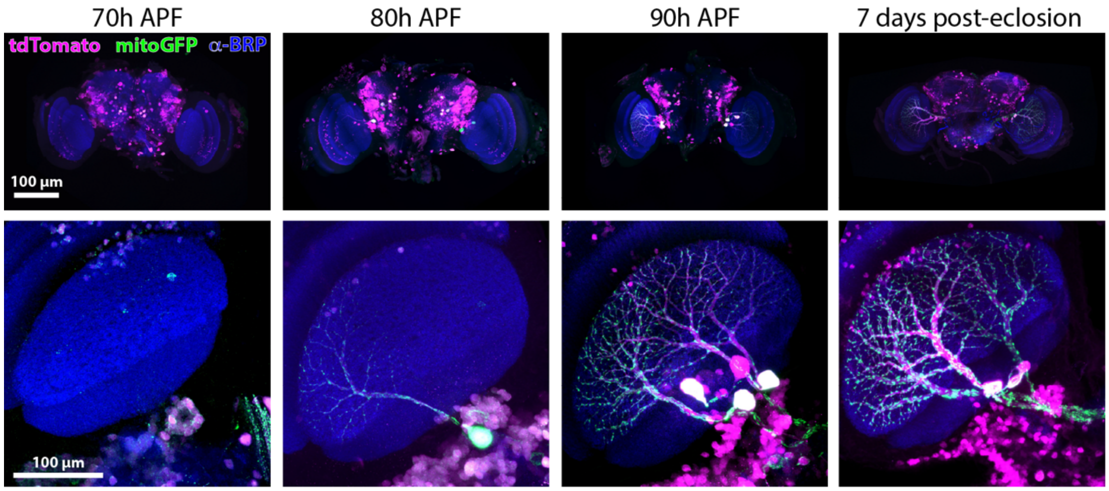


**Figure S1: The R27B03-GAL4 driver line labels HS neurons in late pupation and adulthood.** Representative images of R27B03-GAL4 expression in whole *Drosophila* brains (top row) and the lobula plate of the optic lobe (bottom row) at 70, 80, or 90 hours after pupal formation (APF) and 7 days after eclosion. Neurons were labelled by GAL4-driven expression of tdTomato (magenta) and mitoGFP (green) and the neuropil was labelled by immunostaining for the synaptic marker BRP (blue). All images are maximum projections of z-stacks acquired using confocal Airyscan imaging.

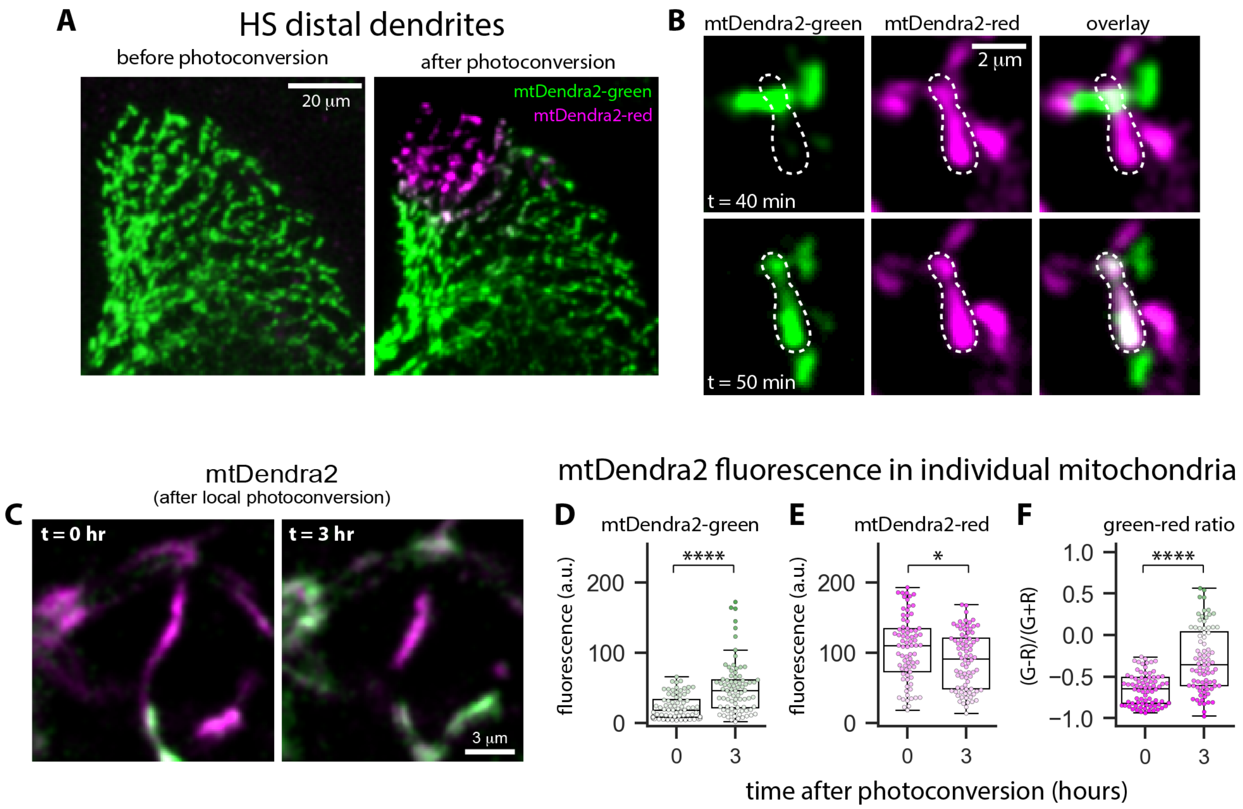


**Figure S2: Mitochondrial fusion and content exchange in HS distal dendrites.** A: mtDendra2-labeled mitochondria in HS distal dendrites before (left image) and after (right image) local photoconversion. Images are max projections of confocal z-stacks. B: Fusion between mitochondria labeled with mtDendra-green and mitoDendra2-red. Images show the mitochondria before (t = 40 min, top row) and after fusion (t = 50 min, bottom row); dashed line indicates a stationary mitochondrion, labelled with mtDendra-red, that receives mtDendra2-green. C: Dendra2-tagged mitochondria in distal HS dendrites immediately after photoconversion (left image) and three hours later (right image). D-F: Quantification of mtDendra2-green fluorescence (D), mtDendra2-red fluorescence (E), and a metric quantifying the green-to-red ratio (F) in individual mitochondria in primary HS dendrites after photoconversion. The green-to-red ratio is given by (G-R)/(G+R), where G is the green fluorescence intensity and R is the red fluorescence intensity. Dots overlaid on the box plots indicate measurements from individual mitochondria; dot color reflects the fluorescence intensity of each mitochondrion (D-E) or the ratio of green-to-red fluorescence within each mitochondrion (F). Asterisks indicate significant differences (Mann Whitney U test; * $p < 0.05$, **** $p < 10^{-5}$); N = 78 mitochondria (t = 0 hours) or 80 mitochondria (t = 3 hours) from 8 neurons.

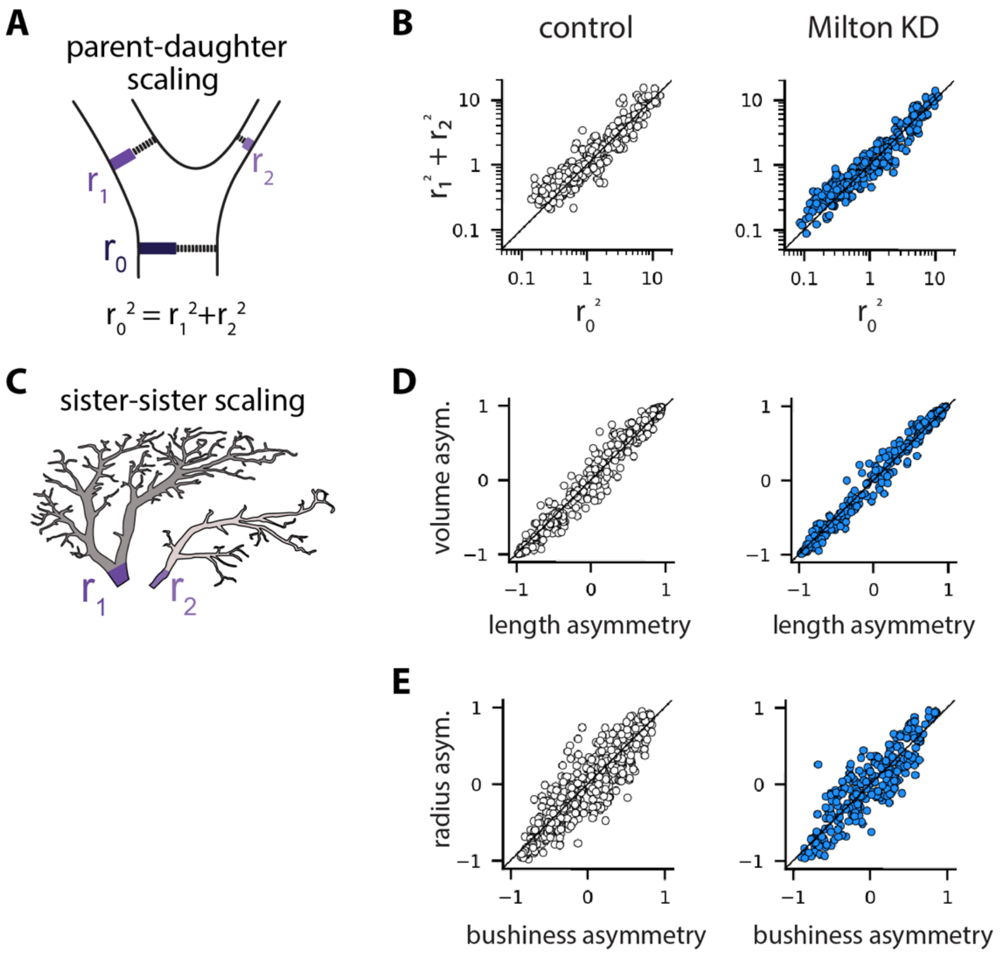


**Figure S3: Milton knockdown has no effect on HS dendrite scaling.** A: Cartoon depicting parent-daughter scaling. Radii at branch points scale according to $r_0^2 = r_1^2 + r_2^2$ , where $r_0$ is the radius of the parent branch and $r_1$ and $r_2$ are the radii of the daughter branches. B: Experimental measurements of parent-daughter scaling in control (open circles, left panel) and Milton KD (blue circles, right panel) dendrites, with $r_0^2$ plotted versus $r_1^2 + r_2^2$. C: Cartoon depicting sister-sister scaling. Sister subtrees scale such that subtree length is proportional to subtree volume and trunk thickness ($r^2$) is proportional to subtree bushiness (length/depth). D-E: Experimental measurements of sister-sister scaling, with subtree length asymmetry plotted versus volume asymmetry (D) and trunk cross-sectional area asymmetry plotted versus subtree bushiness asymmetry (E). N = 438 branch points from 10 dendrites (control) and 256 branch points from 11 dendrites (Milton KD).

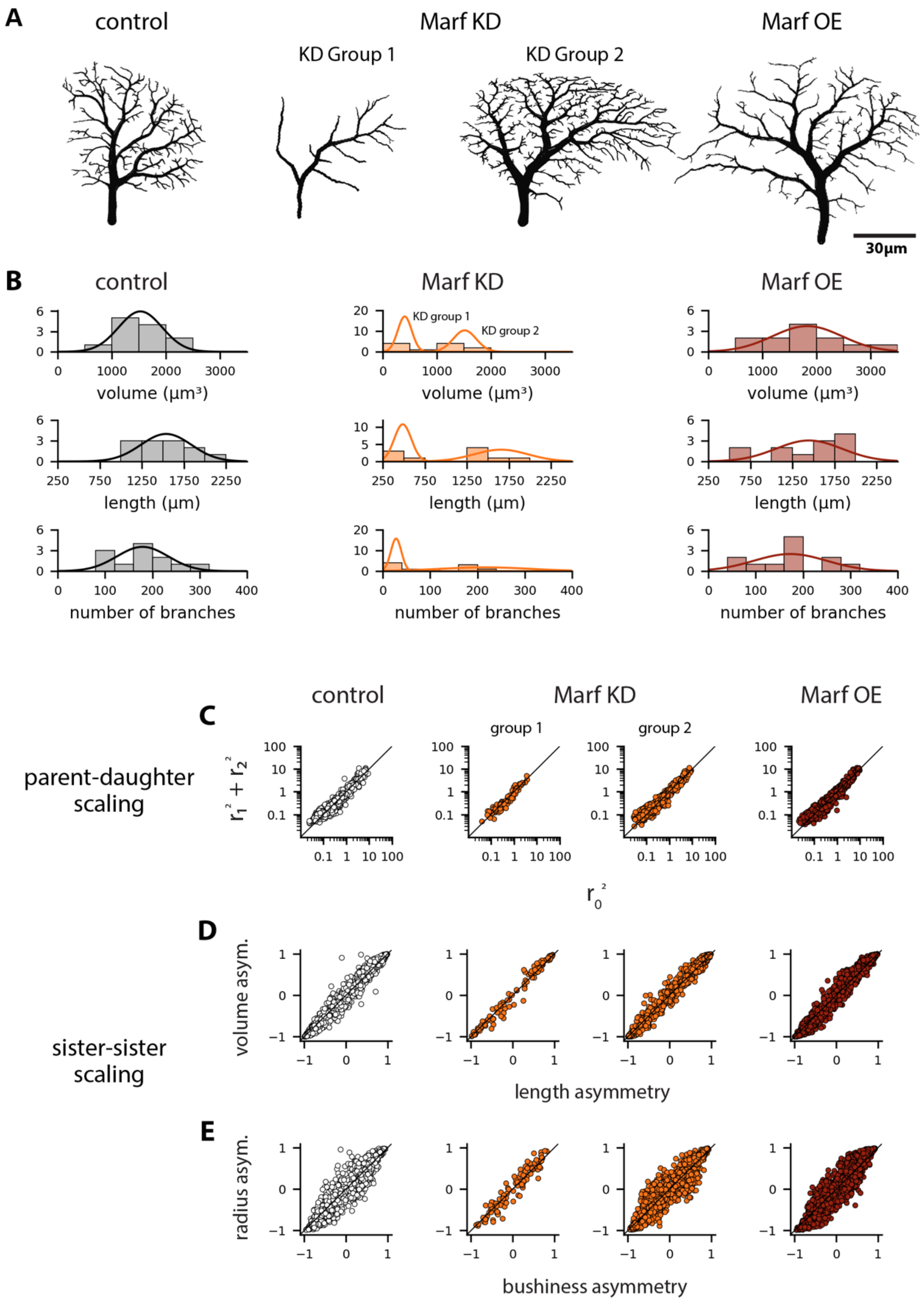
A
control
Marf KD
KD Group 1
KD Group 2
Marf OE
30µm
B
control
Marf KD
Marf OE
KD group 1
KD group 2
volume (µm³)
length (µm)
number of branches
C
control
Marf KD
Marf OE
group 1
group 2
parent-daughter scaling
$r_1^2 + r_2^2$
$r_0^2$
D
sister-sister scaling
volume asym.
length asymmetry
E
radius asym.
bushiness asymmetry

**Figure S4: Marf knockdown has a variable effect on HS dendrite size.** A: Representative reconstructions of HS dendrites from control, Marf knockdown, and Marf overexpression samples. The two Marf knockdown dendrites illustrate the range of dendrite morphologies in the knockdown samples. B: Histrograms showing distributions of dendrite volumes, lengths, and number of branches for control (gray), Marf knockdown (orange), and Marf overexpression (maroon) samples. Gaussian curves overlaid on the histograms were defined by Gaussian mixture models; control and Marf overexpression distributions were best fit by a single Guassian, whereas Marf knockdown distributions were best fit by two Gaussians. C: Parent-daughter scaling in control (open circles, left panel), Marf KD (orange, center panels), and Marf OE (maroon, right panel) dendrites, with $r_0^2$ plotted versus $r_1^2 + r_2^2$. D-E: Sister-sister scaling, with subtree length asymmetry plotted versus volume asymmetry (D) and trunk cross-sectional area asymmetry plotted versus subtree bushiness asymmetry (E). Marf KD samples were divided into two groups based on the bimodal distributions show in B. N = 1564 branch points from 12 dendrites (control), 101 branch points from 5 dendrites (Marf KD group 1), 970 branch points from 6 dendrites (Marf KD group 2), and 1502 branch points from 12 dendrite (Marf OE).

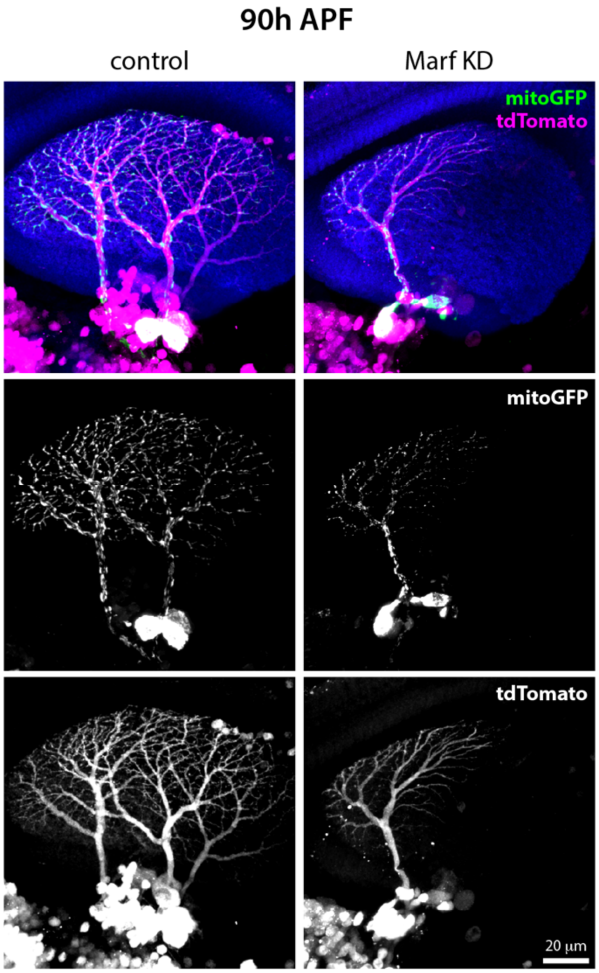


**Figure S5: Marf knockdown reduces HS cell count prior to eclosion.** Representative images of HS dendrites in control (left) and Marf knockdown samples (right) at 90h APF. Neurons were labelled by GAL4-driven expression of tdTomato (magenta) and mitoGFP (green) and the neuropil was labelled by immunostaining for the synaptic marker BRP (blue). Images are maximum projections of z-stacks acquired using confocal Airyscan imaging.

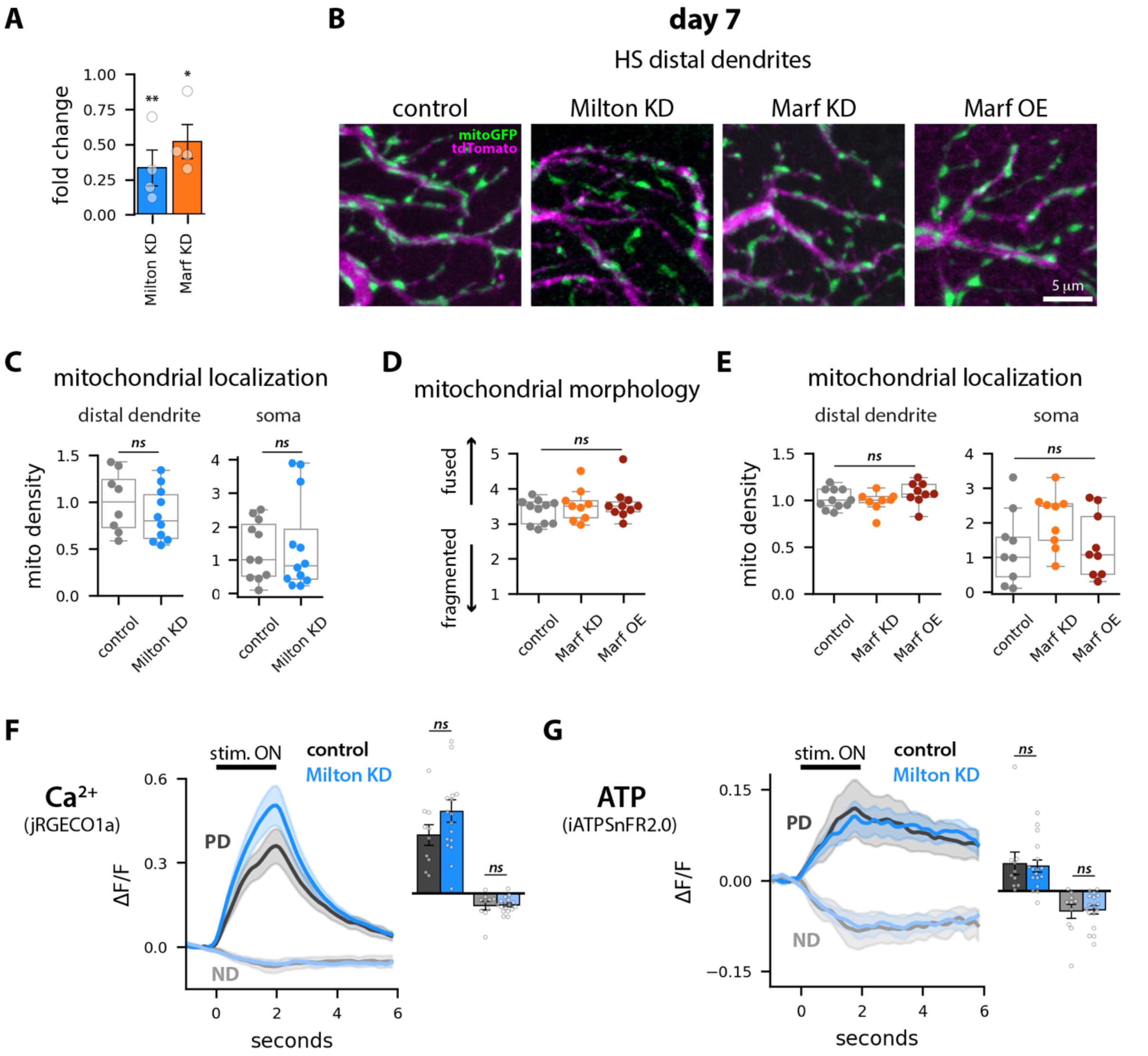


**Figure S6: Adult-restricted Milton knockdown has no effect on mitochondrial localization or visual stimulus-driven ATP responses in young flies.** A: Fold-change in Milton and Marf transcript levels 7 days after pan-neuronal, adult-restricted (with Gal80$^{ts}$) expression of shRNAi. Asterisks indicate significant differences from 1 (* $p < 0.05$; ** $p < 0.01$, one sample T test). B: Representative images of distal HS dendrites 7 days after induction of Milton or Marf knockdown or Marf overexpression. Neurons were labelled with mitoGFP (green) and tdTomato (magenta). Images are maximum projections of confocal z-stacks acquired using Airscan imaging. C: Mitochondrial volume densities, normalized to the median of the control, in distal dendrites (left panel) and somas (right panel) in control (gray) and Milton knockdown (blue) samples. Dots overlaid on the box plots indicate average measurements for individual flies in the distal dendrites (N = 8-9 flies per genotype) or measurements in individual somas (N = 11-12 somas per genotype). D: Quantification of mitochondrial morphology in distal HS dendrites in control (gray), Marf knockdown (orange) and Marf overexpression (maroon) samples. Dots overlaid on boxplots indicate average measurements for individual flies (N = 9-11 flies per genotype). E: Normalized mitochondrial volume densities in distal dendrites (left panel) and somas (right panel). Dots overlaid on the boxplots indicate average measurements for individual flies in the distal dendrites (N = 8-11 flies per genotype) and measurements in individual somas (N = 9 somas per genotype). F-G: Average calcium (jRGECO1a, F) and ATP (iATPSnFR2.0, G) responses to wide field square wave gratings moving in the preferred direction (PD) or null direction (ND) in distal dendrites of control (gray) and Milton knockdown (blue) samples at day 7. Insets show average response amplitudes. Shading on the line plots and error bars on the bar plots indicate the standard error of the mean; dots overlaid on the bar plots are average values for individual flies (N = 10-16 flies per genotype). There are no significant differences among sample groups (Mann Whitney U test, $p>0.05$).
